# Heat ‘n Beat: A universal high-throughput end-to-end proteomics sample processing platform in under an hour

**DOI:** 10.1101/2023.09.27.559846

**Authors:** Dylan Xavier, Natasha Lucas, Steven G Williams, Jennifer M. S. Koh, Keith Ashman, Clare Loudon, Roger Reddel, Peter G. Hains, Phillip J. Robinson

**Author notes:** These authors contributed equally (Dylan Xavier, Natasha Lucas). Sangui Bio, PO BOX 4054, Royal North Shore Hospital, St Leonards, NSW 2065. Sciex, 2 Gilda Ct, MULGRAVE Victoria 3170 (Keith Ashman).

## Abstract

Proteomic analysis by mass spectrometry (MS) of small (≤2 mg) solid tissue samples from diverse formats requires high throughput and comprehensive proteome coverage. We developed a near universal, rapid and robust protocol for sample preparation, suitable for high-throughput projects that encompass most cell or tissue types. This end-to-end workflow extends from original sample to loading the mass spectrometer and is centred on a one tube homogenisation and digestion method called Heat ‘n Beat (HnB). It is applicable to most tissues, regardless of how they were fixed or embedded. Sample preparation was divided to separate challenges. The initial sample washing, and final peptide clean-up steps were adapted to three tissue sources: fresh frozen (FF), optimal cutting temperature (OCT) compound embedded (FF-OCT), and formalin-fixed paraffin-embedded (FFPE). Thirdly, for core processing, tissue disruption and lysis were decreased to a 7 min heat and homogenisation treatment, and reduction, alkylation and proteolysis were optimised into a single step. The refinements produced near doubled peptide yield, delivered consistently high digestion efficiency of 85-90%, and required only 38 minutes for core processing in a single tube, with total processing time being 53-63 minutes. The robustness of HnB was demonstrated on six organ types, a cell line and a cancer biopsy. Its suitability for high throughput applications was demonstrated on a set of 1,171 FF-OCT human cancer biopsies, which were processed for end-to-end completion in 92 hours, producing highly consistent peptide yield and quality for over 3,513 MS runs.

**Graphical Abstract:** 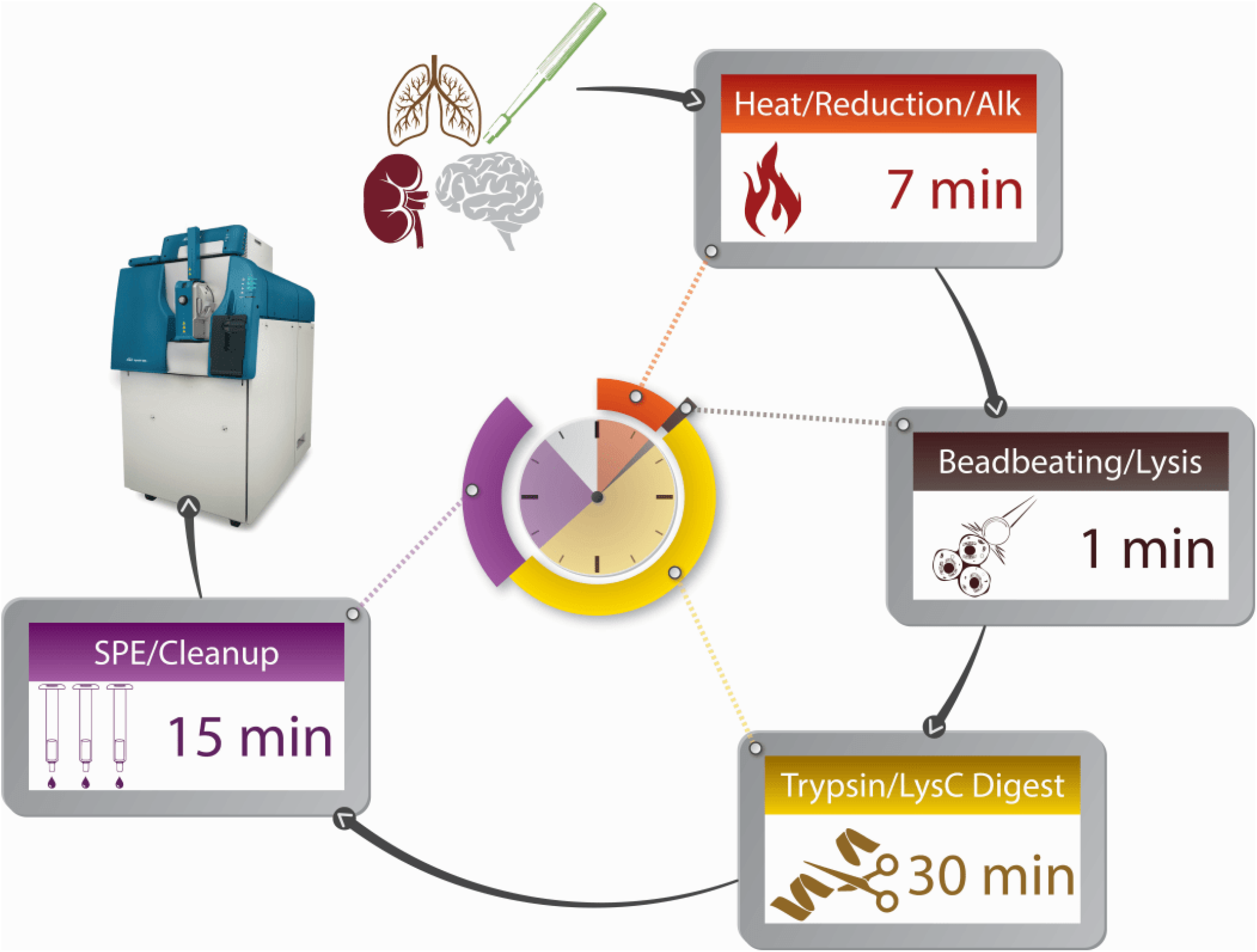

## Introduction

Sample preparation for proteomics by mass spectrometry (MS) has four basic stages after collection that are traditionally performed sequentially: 1) Initial removal of external contaminants (e.g. the embedding material, optimal cutting temperature (OCT) polymer or paraffin, by extensive washing); 2) Tissue disruption and cell lysis by homogenisation or other approaches; 3) Chemical modifications of sensitive amino acid residues and subsequent proteolytic digestion, and 4) Clean-up of the resultant peptides by solid phase extraction (SPE) with associated peptide quantification for loading consistent amounts of peptide on the mass spectrometers. Proteomic workflows need to be streamlined and simplified to allow upscaling of research projects in all fields and to advance clinical applications.

Several studies have described sample preparation methods for MS that enable high-throughput workflows, unfortunately most are typically developed for easy-to-lyse samples like mammalian cell lines or biofluids such as cerebral spinal fluid, plasma, or urine ^1–4^. Solid tissue punches or sections and difficult-to-homogenise tissues like muscle and formalin-fixed tissues present greater solubilisation and protein extraction challenges ^5–7^. The latter can often be overcome with the use of pressure cycling technology (PCT) in a Barocycler instrument ^8^. However, current methods do not support true high-throughput sample preparation and are not sufficiently flexible to accommodate large variations in the types of tissues sources.

We previously reported a sample preparation method called Accelerated Barocycler Lysis and Digestion (ABLE) ^9^, that optimised PCT to prepare small tissue biopsies (≤2 mg) for MS analysis in four hours (end-to-end time, defined as from cut sample to ready to load the mass spectrometer). ABLE substantially reduced the time and cost of tissue sample processing compared to the earlier published PCT-SWATH methods. However, ABLE utilises a multi-step procedure that requires time-consuming steps such as uncapping and recapping of tubes between steps, potentially introducing variability due to sample loss between tubes and steps. ABLE was not optimised to suit formalin-fixed paraffin-embedded (FFPE) tissue.

To date, no published method provides an end-to-end solution for sample preparation across the diversity of proteomic sample types, particularly in the case of human tissues that have been processed routinely as FFPE blocks or fresh frozen (FF) specimens embedded in OCT for cryosectioning. We aimed to develop a universal ‘one pot’ method that can be used with multiple tissue types and across numerous preservation techniques, which is fast, robust and has automation potential. The method should also be reproducible and high-throughput. Many proteomic workflows include an overnight digestion, which is unsuitable for processing in a clinically relevant timeframe ^10–16^. Another study compared the performance of the 16 most commonly used methods have been compared ^17^ (see also **Supplementary Table S1**). Here, we show that it is possible for the principal “core processing” (tissue disruption and tryptic digestion, **Figure 1**) to be performed by a single common strategy for essentially all sample sources. Following this, we show how the two main embedding materials can be dealt with using fast front- and back-end approaches, resulting in a true end-to-end method.

**Figure 1:**
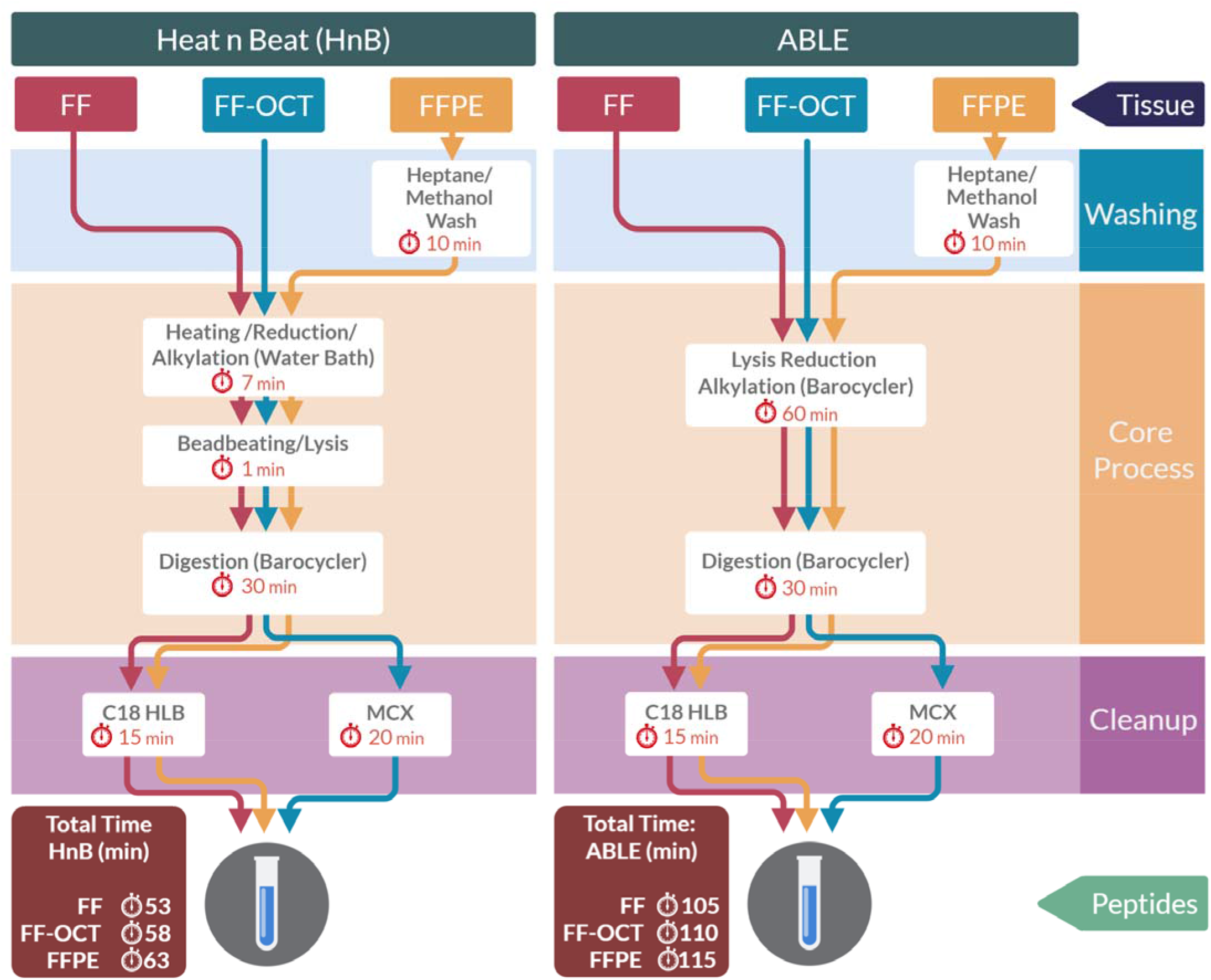
Summary of Heat ‘n Beat (HnB) protocol vs ABLE. End-to-end sample preparation workflow when processing fresh frozen (FF), optimal cutting temperature (OCT) compound embedded (FF-OCT), and formalin-fixed paraffin-embedded (FFPE) tissue samples prepared with either the HnB or the ABLE method. The steps (right) are divided into sections for washing and peptide clean-up flanking the central core processing.

To develop the common core process, we used ABLE as a starting point and progressively modified the emergent method by identifying and addressing challenges, bottlenecks and limitations with existing tissue disruption and tryptic digestion methods. Key amendments were made to the tissue disruption and protein digestion steps consolidated into a one tube core processing method we call the Heat n Beat (HnB) protocol that reduced the time required to process samples to under an hour (typically 38 min). To allow for a broader range of sample inputs to HnB, we explored two key approaches for embedding reversal and two methods for SPE, that are required to accelerate processing of FF-OCT or FFPE samples. We developed methods to reduce the time required for embedding reversal to a few minutes (FFPE) or not at all (FF-OCT). To validate the performance of HnB in a high-throughput scenario, a large human biopsy cohort of 1,171 FF-OCT samples was used, for which total end-to-end sample preparation took 92 hours and which resulted in excellent peptide yields, consistently high peptide quality and protein group identification. Our HnB end-to-end method demonstrates the feasibility of analysing large diverse tissue cohorts, to produce consistently reliable results, in a clinically relevant timeframe.

## Materials/Methods

### Tissue Processing

Whole organs were harvested from rats, with approval from the Animal Care and Ethics Committee for the Children’s Medical Research Institute, Sydney, Australia (project number C116). The following seven rat organs with a range of expected homogenisation difficulty were used: brain, testis, spleen, kidney, lung, liver, heart and leg muscle (gastrocnemius). They were obtained from adult male rats (>300 gm body weight) which were euthanised by cervical dislocation. Three proteomic sample preparation methods were applied to these samples, each from the same starting tissue to facilitate direct comparison between methods: fresh frozen tissue (FF) on dry ice, fresh frozen and embedded in OCT compound (FF-OCT), or formalin-fixed and paraffin embedded (FFPE). For the FF and FF-OCT samples, the left side of the brain, half of the right and left testes, half of the right and left leg muscle, half of the liver, half of the right and left kidney, half of the heart, half of the spleen and the right lung were rapidly frozen by immediately placing in a conical tube on dry ice and transferring to a −80_°_C freezer for storage. For FFPE samples, the remaining halves of each organ were immediately immersed in 10% neutral buffered formalin (conical tube) solution and fixed for 24 hrs at 22_°_C.

The formalin-fixed organs were cut into suitable pieces and transferred to labelled tissue cassettes. They were immersed in 70% (v/v) ethanol and processed by automation on an embedding station (Excelsior AS Tissue Processor, ThermoFisher Scientific) at Westmead Institute of Medical Research (Sydney). The automated cycle started with dehydration in 70% (v/v) ethanol for 10 min, then increasing alcohol concentration in five consecutive steps from 75-100% (v/v) at ambient temperature (total time 6.5 hrs). This was followed by three separate incubation steps in fresh lots of xylene at ambient temperature for 3 x 1 hr. Finally, the tissue samples were infused in three changes of paraffin wax at 60_°_C for 80 min. Tissues embedded in paraffin were stored at room temperature (approx. 22_°_C).

Human prostate cancer tissues were collected with informed consent and sourced from the Department of Pathology and Molecular Pathology at the University Hospital Zurich, Switzerland. Men scheduled for radical prostatectomy with clinically localised prostate cancer were selected from the Prostate Cancer Outcomes Cohort (ProCOC) study ^18^. This study was approved by the Cantonal Ethics Committee of Zurich, and each patient provided informed consent (KEK-ZH-No. 2008-0040). The tissue was embedded in OCT compound. The use of prostate tissue samples in this study was approved by the Western Sydney Local Health District Human Research Ethics Committee (AU RED HREC/17/WMEAD/63). For the large cohort study, 1,171 frozen human cancer biopsies were obtained from the Victorian Cancer Biobank (VCB; https://viccancerbiobank.org.au/) and were sectioned and embedded in OCT (FF-OCT).

Samples were cut from the FF specimens (rat organs) using 3 mm biopsy punches. For FF-OCT samples, Sakara Tissue Tek OCT compound was added to the FF tissue and sectioned at 30 µm using a Leica CM1860 UV cryostat. For the FFPE samples, 10 µm sections were cut. The curls were transferred to a Barocycler MicroTube (up to 150 µL volume, from Pressure Biosciences, South Easton, MA, USA). These low protein binding tubes are constructed from fluorinated ethylene propylene for integrity over a wide temperature and pressure range. Finally, all samples were stored in a Barocycler MicroTube at −20_°_C – we refer to this as a ‘one tube’ method as all subsequent steps utilise the same tube.

HEK293T cells (from Dr. Timothy Adams, CSIRO Manufacturing, Parkville, Victoria, Australia) were adapted to grow in suspension in Freestyle 293 Expression medium (Life Technologies Australia, Scorsby, VIC, Australia) supplemented with 200 mg/L G418 (Life Technologies) using a humidified shaker incubator (37°C, 5% CO_2_, 130 rpm). The adapted HEK293T culture was maintained in Erlenmeyer shaker flasks and scaled up in a 20 L WAVE bioreactor (GE Healthcare, Silverwater, NSW, Australia), seeded at an initial working volume of 5 L at a concentration of 0.8 × 10^6^ viable cells/mL. To reduce shear stress, Pluronic F68 (Life Technologies) was added to the culture at 0.2% (w/v) final concentration. At a viable cell density of 3 × 10^6^ cells/mL the culture was scaled up to 20 L and maintained at 37°C with a rocking speed of 25 rpm and rocking angle of 9°. The cells were harvested at _∼_5 × 10^6^ cells/mL by centrifugation (1500 xg, 10 min, 4°C), snap-frozen on liquid nitrogen and stored at −80°C. Frozen cell pellets were scraped using a small spatula into a Barocycler MicroTube.

### Deparaffinisation of FFPE tissue

The final deparaffinisation method for FFPE samples was adapted from Jullig *et al.* ^19^, for the 10 µm FFPE tissue in Barocycler MicroTubes, 120 µL of heptane was added to dissolve the paraffin and the tubes were incubated in an Eppendorf ThermoMixer C at 30°C for 10 min with shaking at 900 rpm. Then 30 µL of methanol was added to the heptane to create a two-phase liquid layer. The upper (heptane) layer was removed and the methanol evaporated under reduced pressure using a GeneVac^®^ EZ-2 plus evaporator. This procedure takes less than 12 minutes.

### Removal of OCT via washing

For comparison with OCT removal post lysis and digestion using MCX, we also performed the following protocol for tissue washing based on the protocol prepared by M. Martinez and K. Shaddox, Vanderbilt-Ingram Cancer Center, published by The National Cancer Institute’s Clinical Proteomic Tumor Analysis Consortium (CPTAC) ^8^. Tissue (in a 1.5 mL Eppendorf tube), was washed with 1 mL 70% (v/v) EtOH, vortexed for 30 s, centrifuged briefly and the supernatant discarded. This was repeated with 1 mL of 100% H_2_O. Next 1 mL of 70% (v/v) EtOH was added and incubated at room temperature for 5 min, then centrifuged 20,000 xg for 2 min at 20_⁰_C, and the supernatant discarded. This was repeated once with 70% (v/v) EtOH, the twice with 85% (v/v) EtOH, and twice with 100% (v/v) EtOH. This procedure takes approximately 1 hr.

### Accelerated Barocycler Lysis and Extraction (ABLE)

We previously reported an Accelerated Barocycler Lysis and Extraction (ABLE) protocol for proteomics sample preparation which serves as the baseline comparator for this investigation ^9^. Briefly, tissue lysis, reduction and alkylation were carried out simultaneously using 30 µL of 1% (w/v) sodium deoxycholate (SDC), 5% (v/v) *N*-propanol, 100 mM triethylammonium bicarbonate, 10 mM tris(2-carboxyethyl)phosphine (TCEP) and 40 mM iodoacetamide (IOA) in a Barocycler 2320EXT instrument at 45 kpsi, for 60 cycles (50 s high pressure, 10 s atmospheric pressure) at 56°C. Following this, 120 µL of Rapid Digestion Buffer (Promega, VA1061) and 1 µg of Rapid Trypsin/Lys-C (Promega, VA1061) was added to each sample. Digestion was carried out in the Barocycler using 30 cycles of 50 s at 45 kpsi and 10 s at atmospheric pressure and at 70°C. Samples were acidified with 5 µL of formic acid, to precipitate the SDC which was removed by centrifugation (15 min, 18,000 xg, 4°C). The supernatant was transferred to a new tube and cleaned up using solid phase extraction (SPE) on HLB resin. The eluent was evaporated to dryness and samples were resuspended in 0.5% (v/v) formic acid. Note that in the initial ABLE protocol, OCT was removed from the relevant samples prior to lysis, by a method based on that of Zhang *et al.* ^20^, which takes about an hour. However, for the present study, OCT was not removed prior to lysis, instead a novel strategy was deployed for HnB (below).

### Heat ‘n Beat (HnB)

To develop and validate the final central HnB one tube homogenisation and digestion protocol and to determine its efficacy, the seven different rat tissues (kidney, muscle, liver, brain, testis and lung, spleen) were prepared in three ways (FF punches, or FF-OCT and FFPE sections) and each was processed in parallel using ABLE or HnB. To support mass spectrometry the paraffin was removed from FFPE tissue using the rapid heptane-methanol liquid-phase described above, while OCT was not removed until the final SPE stage (**Figure 1**). As cell lines are usually simply frozen rather than being fixed and/or embedded, human HEK293T cells were grown as previously described ^9,21^ and FF cell pellets were also prepared using both ABLE and HnB. Prostate samples were prepared as described for FF-OCT above for 30 µm sectioning and prepared using ABLE and HnB, but were washed prior to lysis or cleaned up using MCX SPE (below).

For HnB sample preparation, the core processing steps were the same for all samples in this investigation (**Figure 1**). After the relevant washing steps, proteins were denatured, reduced and alkylated in their original Barocycler MicroTubes using 11 µL of 5% (w/v) sodium deoxycholate (SDC), 100 mM triethylammonium bicarbonate, 4.2 mM tris(2-carboxyethyl)phosphine (TCEP) and 16.75 mM iodoacetamide (IOA) and incubated in a water bath at 95°C for 7 min. Following this, 50 µL of rapid digestion buffer (Promega), 1 µg of trypsin/Lys-C (Rapid Trypsin, Promega^©^), and 1 unit of Benzonase nuclease (Sigma, St. Louis, MO). Then twelve 1 mm TriplePure Zirconium beads (Benchmark Scientific, Sayreville, NJ, USA, D1132-10TP/250 g) were added to support bead-beating. Samples were capped with 50 µL MicroCaps (Pressure Biosciences, South Easton, MA, USA) and placed in a Beadbug 3 instrument (Benchmark Scientific, Sayreville, NJ, USA, D1030), which was modified with an in-house custom adaptor to hold Barocycler cartridges housing 16 Barocycler MicroTubes together, and horizontally shaken in the bead beater at 3,800 rpm for 1 min. Tubes were transferred to a bench top Barocycler 2320EXT using 30 cycles of 50 s at 45 kpsi and 10 s at atmospheric pressure and at 56°C (total time 30 min), where lysis and digestion were completed. Samples were acidified with 5 µL of formic acid to precipitate the SDC, then pelleted by centrifugation (15 min, 18,000 x g, 4°C). The supernatant containing the peptides was transferred to an Eppendorf tube and cleaned up using the appropriate one of the two SPE methods below (**Figure 1**).

### Solid phase extraction (SPE)

Two forms of SPE were used to clean-up the final pool of peptides for MS analysis. Peptides from FF punches, FFPE sections and HEK293T cell line samples were cleaned and concentrated after core processing using the following Hydrophilic-Lipophilic Balance (HLB) protocol. However, peptides from FF-OCT embedded fresh frozen sections for both HnB and ABLE preparation were cleaned and concentrated after core processing using the following Mixed Cation Exchange (MCX) protocol, designed to remove the polymers from the OCT.

### Hydrophilic-Lipophilic Balance (HLB)

Oasis HLB is a polymeric reversed phase sorbent made from a copolymer of the hydrophilic N-vinylpyrrolidone and the lipophilic divinylbenzene and which has a performance similar to traditional silica-based SPE particles like octadecyl (C18) resin, except with a 3X higher hydrophobic retention capacity. Oasis HLB SPE cartridges (1 cc, 30 mg, Waters, Rydalmere, NSW, Australia) were conditioned using 1 mL 90% (v/v) acetonitrile, 0.5% (v/v) formic acid and equilibrated with 1 mL of 0.5% (v/v) formic acid. Samples were diluted up to 1 mL using 0.5% (v/v) formic acid and loaded onto the cartridges. SPE columns were then washed twice with 1 mL of 0.1% (v/v) formic acid before being eluted with 200 µL of 70% (v/v) acetonitrile, 0.5% (v/v) formic acid followed by 100 µL of 90% (v/v) acetonitrile, 0.5% (v/v) formic acid. The eluent was evaporated to dryness and samples were resuspended in 0.5% (v/v) formic acid and the concentration determined as described below. This procedure typically takes 15 minutes.

### Mixed Cation Exchange (MCX)

Oasis MCX is a mixed-mode polymeric sorbent with both reversed phase and cation-exchange functionalities, with the strong cation-exchange sulfonic acid groups in addition to the above Oasis HLB sorbent. These properties allow for maximum comparison with the above HLB method since the base resin is the same. Oasis MCX SPE cartridges (1 cc, 30 mg, Waters) were conditioned using 1 mL 100% methanol and equilibrated with 1 mL of 2% (v/v) formic acid. Samples were diluted up to 1 mL using 2% (v/v) formic acid and loaded onto the cartridges. To remove the OCT and other contaminants, the cartridges were then washed in succession with 1 mL of 2% (v/v) formic acid, two washes of 1 mL of 60% (v/v) acetonitrile, 2% (v/v) formic acid, 1 mL 5% (v/v) acetonitrile, 1 M ammonium formate, 2% (v/v) formic acid, 1 mL 2% (v/v) formic acid and two washes of 1 mL ultrapure water. Peptide samples were then eluted with 500 µL of 30% (v/v) acetonitrile, 5% (v/v) ammonium hydroxide. The eluent was evaporated to dryness and samples were resuspended in 0.1% (v/v) formic acid and the concentration was determined as described below. This procedure takes 20 minutes.

### Peptide quantitation

For all approaches, the final peptide yield was quantified from the concentration of the peptide solution determined by UV absorption using A280 nm with an Implen nanophotometer N60 (LabGear, Brisbane, QLD, Australia).

### Liquid chromatography-mass spectrometry (LC-MS/MS) methods

#### Information Dependent Acquisition (IDA)

An Eksigent nanoLC 425 HPLC operating in microflow mode, coupled online to a 6600 Triple TOF (Sciex) was used for the analyses. The peptide digests (2 µg) were injected onto a C18 trap column (SGE TRAPCOL C18 G203 300 µm x 100 mm) and desalted for 5 min at 10 µL/min with solvent A (0.1% [v/v] formic acid). The trap column was switched in-line with a reversed phase capillary column (SGE C18 G203 250 mm × 300 µm, internal diameter 3 µm, 200 Å), maintained at a temperature of 40°C. The flow rate was 5 µL/min. The gradient started at 2% solvent B (99.9% [v/v] acetonitrile, 0.1% [v/v] formic acid) and increased to 10% over 5 min. This was followed by an increase of solvent B to 25% over 60 min, then a further increase to 40% for 5 min. The column was washed with a 4 min linear gradient to 95% solvent B held for 5 min, followed by a 9 min column equilibration step with 98% solvent A. The HPLC eluent was analysed using the Triple TOF 6600 system (Sciex) equipped with a DuoSpray source and 50 µm internal diameter electrode, controlled by Analyst 1.7.1 software (AB Sciex 2015). The following parameters were used: 5500 V ion spray voltage; 25 nitrogen curtain gas; 100°C TEM, 20 source gas 1, 20 source gas 2. The 90 min information dependent acquisition (IDA) consisted of a survey scan of 200 ms (TOF-MS) in the range 350–1250 m/z to collect the MS1 spectra and the top 40 precursor ions with charge states from +2 to +5 were selected for subsequent fragmentation with an accumulation time of 50 ms per MS/MS experiment for a total cycle time of 2.3 s and MS/MS spectra were acquired in the range 100–2000 m/z.

#### Data-Independent Acquisition (DIA/SWATH)

For DIA/SWATH acquisition peptide spectra were acquired with the LC MS/MS method as described for IDA acquisition, using 100 variable windows. The parameters were: lower m/z limit 350; upper m/z limit 1250; window overlap (Da) 1.0; CES was set at 5 for the smaller windows, then 8 for larger windows; and 10 for the largest windows. MS2 spectra were collected in the range of m/z 100 to 2000 for 30_ms in high resolution mode and the resulting total cycle time was 3.2 s.

### Data Processing

IDA spectra were searched using ProteinPilot™ Software v5.0 (Sciex) using the Paragon algorithm with the following parameters: Sample Type: Identification; Cys Alkylation: Iodoacetamide; Digestion: Trypsin; Instrument: TripleTOF 6600; Database: Uniprot Rat (37,596 entries) for both fresh frozen and FFPE rat tissues. Thorough ID and False Discovery Rate (FDR) Analysis were selected, and the FDR was set at 1%.

For processing of DIA/SWATH data using Data-Independent Acquisition by Neural Networks (DIA-NN) software (version 1.8) ^22^, we created an *in silico* spectral library for the canonical rat proteome (Uniprot Release 2019_05; 8,068 sequences), plus retention time peptides. Data from 60 SWATH .wiff files were used to create a spectral library in DIA-NN. The spectral reference library (SRL) contained 4217 protein isoforms, 4763 protein groups and 54666 precursors in 44219 elution groups. The 60 files were processed in DIA-NN using this SRL and the following settings: Precursor ion generation using: deep learning-based spectra and RTs prediction, protease trypsin/P with 1 missed cleavage, *N*-terminus M-excision and C-carbamidomethylation and no variable mods; peptide length range 7-30; Precursor m/z range 400-1250; fragment ion m/z range 100-2000; output filtered at 0.01 FDR; Scan window radius set to 10; mass accuracy fixed to 2e-05 (MS2) and 1.2e-05 (MS1); interference removal disabled. Final output data were log2-transformed and normalised using LOESS in InfernoRDN ^23^.

### Data Availability

The raw mass spectrometry proteomic data have been deposited in the ProteomeXchange Consortium via the PRIDE partner repository ^24^ with the dataset identifier PXD045405.

## Results and Discussion

We systematically optimised the ABLE proteomic sample preparation protocol ^9^ to deliver a universal method for high-throughput end-to-end sample preparation prior to MS that is suitable for most common tissue types, regardless of the main embedding or fixation methods. Two of the three main sample sources used here required different washing and embedding removal methods and different peptide clean-up procedures, but all types of solid samples share the central core processing HnB pipeline (**Figure 1**). In the first two sections we report improvements in the sample washing and embedding removal procedures at either end of the workflow, streamlining and introducing compatible changes.

### Optimisation FFPE tissue washing

Crosslinking of FFPE tissue by formaldehyde preserves the architecture of tissue, mainly by forming methylene bridges between basic amino acids ^25,25^, while the paraffin embeds the tissue for easier sectioning at room temperature. Good coverage of the proteome in FFPE tissue requires deparaffinisation and removal of crosslinks to reverse the chemical modifications masking peptide identification, notably lysine methylation ^26^. However, FFPE sample processing produces only relatively minor modifications, affecting 2–6% of peptide-spectrum matches ^12,26^. The most common processes for reversing the embedding of FFPE samples is a combination of deparaffinisation, rehydration of the tissue, and heating for antigen retrieval, but most protocols require 25-110 min and involve transfer of the tissue samples between multiple tubes, which is a significant source of sample loss ^12,19,27–30^. We initially used rat kidney FFPE tissue to test a range of optimisations without any tube transfers until the final SPE clean-up step.

Heptane or hexane are among the most commonly used solvents to remove wax from FFPE samples ^19,31^. Heptane is a safer and less toxic substitute for aromatic or chlorinated solvents like xylene. Sub-X is an odorless aliphatic hydrocarbon xylene substitute which offers another alternative ^32^, but in our hands Sub-X did not fully dissolve the wax, even after 1 hr of incubation and shaking. Therefore, heptane deparaffinisation was used. After heptane, there are two common methods for tissue rehydration. Using one method, heptane is removed by step-wise ethanol rehydration with 100% EtOH, 85-90% EtOH, 70% EtOH (added to the tissue and incubated with mixing for 5 min) followed by drying the tissue. Alternatively, a liquid-phase extraction eliminates the time-consuming rehydration steps (**Figure 2a**), wherein methanol is added to the heptane, the upper (heptane) layer is removed, and the methanol is evaporated ^33^. There was no significant difference between the two methods in terms of peptide and protein identification (ID) (**Figure 2b-c**). The heptane methanol extraction was selected, as it yielded cleaner tissue (determined by visual inspection showing no paraffin remaining in the tube) in the shortest time (10 min) with the fewest number of steps.

**Figure 2:**
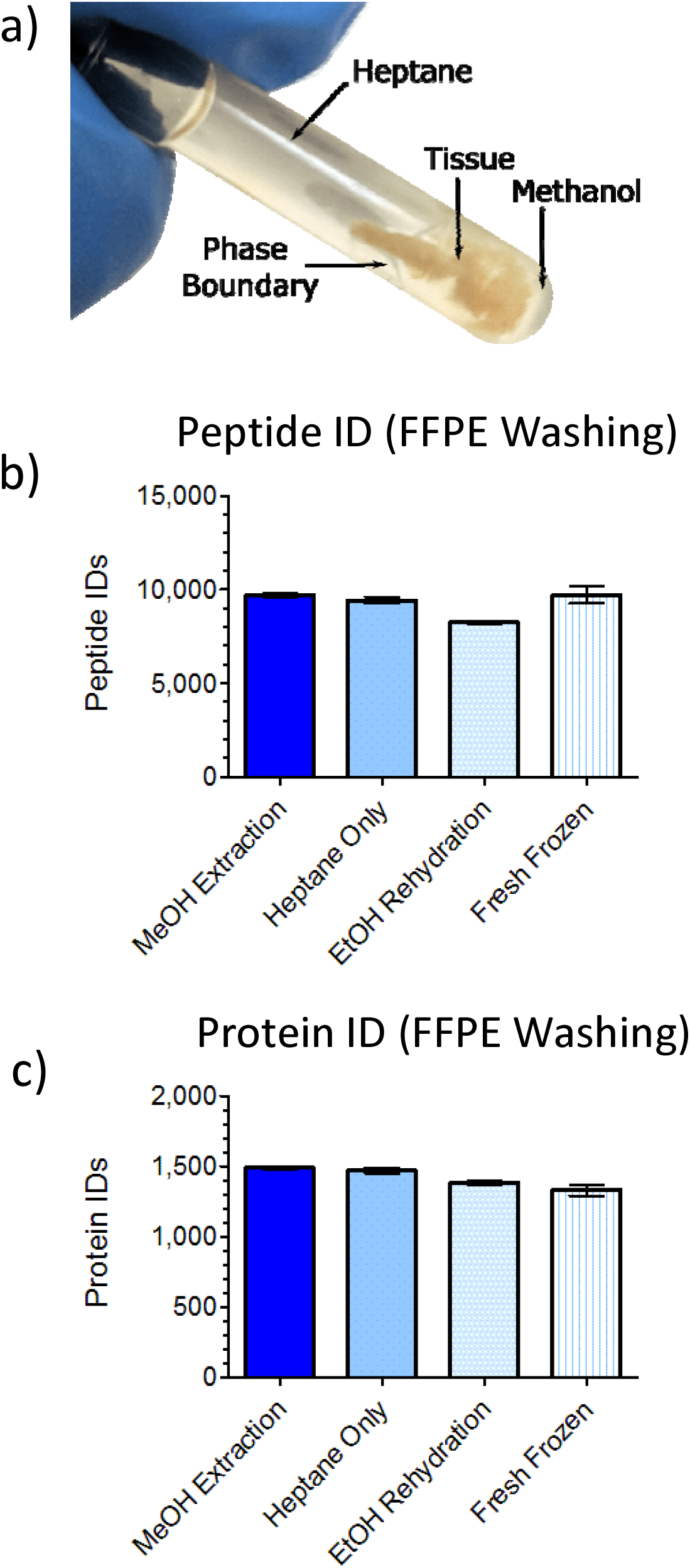
Comparison of FFPE tissue washing methods. **a)** Rat kidney FFPE sections were incubated in heptane to remove wax and then either rehydrated using a liquid/liquid extraction in MeOH, no rehydration (heptane only), a stepwise rehydration with EtOH. The phase boundary between the MeOH and heptane is demonstrated **b-c**) The number of peptide (b) or protein (c) identifications observed from the rat kidney FFPE samples using 3 different washing procedures were compared to unfixed fresh frozen samples (FF). Data are means ± SEM for n = 4 biological replicates.

Following deparaffinisation and rehydration, antigen retrieval is typically performed at high temperature ^27^. Most methods utilise a pre-incubation step at 95-100°C for 30-90 min ^29,30,32^. Two pre-incubation times of 30 or 60 min at 100°C were compared with no pre-incubation, but no significant differences in identifications were observed (**Supplementary Figure S1a**). In contrast, 30-60 min heating was detrimental to the sample and hindered the alkylation step, as evidenced by a major reduction in cysteine carbamidomethylation (presumably due to a reaction with the iodoacetamide that is used to block cysteine from oxidation, **Supplementary Figure S1b**). Therefore, long heating times were avoided in this study. The final 10 minute protocol, resulted in a simple single-tube washing method, ready for core-processing. Three pressure cycling options available on the Barocycler 2320EXT were evaluated. The instrument allows for the pressure to be kept constant for a fixed amount of time, the traditional on/off cycling, or a sine wave pressure cycling which oscillates the pressure around a set value (40 kpsi ± 5 kpsi). There was no advantage for any cycling type and the traditional on/off cycling method was retained (**Supplementary Figure S1c**).

### Eliminating washing of OCT-embedded tissues with MCX

OCT is removed by multiple washing steps, but in our hands, existing protocols often resulted in incomplete removal, as determined by polymer contamination of the MS runs. One of the most rigorous methods for OCT washing is the protocol prepared by M. Martinez and K. Shaddox, Vanderbilt-Ingram Cancer Center, published by The National Cancer Institute’s Clinical Proteomic Tumor Analysis Consortium (CPTAC) ^8^. This is typically performed in a 1.5 mL Eppendorf tube, and in our case the sample is subsequently transfered to a relatively low volume (150 µL) Barocycler MicroTube. Successful washing becomes problematic with softened tissue, and a reduced dead volume is available for mixing space above the samples in small tubes, and the overall process is a source of significant sample loss. In the ABLE protocol, FF-OCT sample washing in low volume Barocycler MicroTubes removed sufficient OCT (success was defined as not interfering with MS analysis) only when using small tissue punches with minimal residual OCT remaining after trimming ^9^. To eliminate the tissue washing step, OCT was removed at the end-stage using Oasis MCX microcolumns after all core processing steps. This mixed-mode resin allows for binding peptides to the reversed phase and strong cation exchange sorbents, with the OCT being retained on the reversed phase resin. Washing the column with 2% formic acid removed additional salts while peptide binding to the ion-exchange resin was improved. The OCT was then eluted from the reversed phase bed with high organic (60% ACN), with the peptides retained on the ion-exchange bed. The bound peptides were washed with formic acid and water and then finally eluted with ammonium hydroxide. Notably, an unidentified contaminant was observed in the eluate that interfered with peptide quantitation by UV spectrophotometry. This was removed by an ammonium formate wash prior to peptide elution. The method provided a clean peptide sample with only a 5 min increase in SPE handling time over the more traditional HLB SPE method.

The optimised MCX method for SPE was compared with the CPTAC washing method on FF-OCT rat kidney and human prostate samples (30 µm sections placed in Barocycler MicroTubes). Sections were washed using the CPTAC protocol, processed using ABLE and cleaned up by the reversed phase HLB SPE, while other sections were processed with ABLE (no prior tissue washing; OCT was present during core processing steps) and clean-up was by the developed MCX SPE method. Both successfully removed the OCT, since no polymer traces were found in the MS spectra. They also resulted in comparable numbers of identified protein groups and peptides (**Figure 3a-b**), but the MCX method significantly increased total peptide yield (*p*-value <0.001) (**Figure 3c**). The presence of OCT during the core processing steps did not interfere with digestion efficiency (**Figure 3d**). It is worth noting that if samples to be analysed contained higher concentrations of OCT than used here, this could require an adjustment of the sorbent bed volume, as saturating OCT has the potential to outcompete peptides. We used a 30 mg sorbent bed, for 150 µL MicroTubes full of OCT, however, trimming of excess OCT from samples prior toning is advisable for some samples. An additional, 60% ACN washing steps can be added to ensure heavy loads of OCT are removed. Overall, the MCX SPE method for end-stage OCT removal saves at least 1 hr, eliminates sample loss due to tube transfers, eliminates the small number of failed washes, and almost doubles the peptide yield.

**Figure 3:**
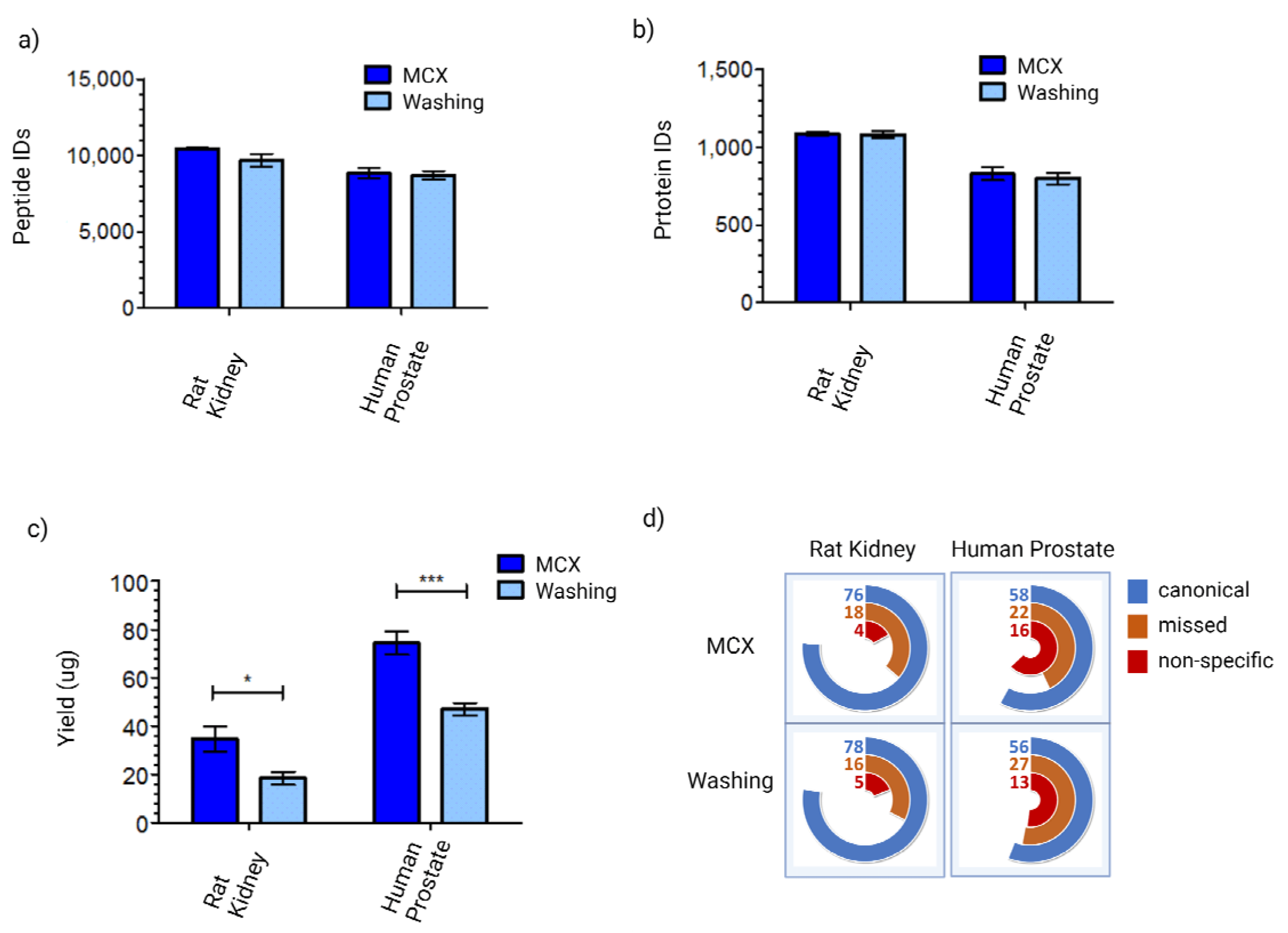
SPE on MCX resin eliminates the need for tissue washing FF-OCT embedded samples. The impact of eliminating OCT removal from FF-OCT embedded samples prior to core sample processing. Rat kidney or human prostate cancer FF-OCT samples (30 µm sections) were washed using the CPTAC washing protocol (washing, dark blue) or washing was omitted (MCX, light blue). All samples were processed using the ABLE protocol and the resultant peptides, with or without OCT being present, and the peptides were cleaned up using MCX SPE. After MS analysis by IDA, the number of **a**) peptides or **b**) protein identifications are shown, along with **c**) the total peptide yield and **d**) digestion efficiency (shown as the percent of canonical trypsin/Lys-C cleaved peptides). Data are means ± SEM for n = 4 biological replicates and p <0.05 (*) or <0.001 (***).

### Development of HnB

Next we focussed on developing a “core processing” method which can be performed by a single common strategy for all three solid tissue sample sources. To develop this, we first used FF rat kidney (collected as tissue punches) as there could be no impact of prior OCT embedding or FFPE fixation. ABLE was used as a starting point and the steps were incrementally modified, with each step being benchmarked against the ABLE protocol, using criteria including: the total number of peptides and protein groups identified and digestion efficiency. The method development is described here using ABLE or HnB with incremental changes to each group (**Figure 4: A-E, or V1-5**). The first step was to simplify core processing into a ‘one tube’ method (**Figure 4a, V1**). ABLE includes two steps in 150 µLtubes tissue lysis, protein solubilisation, reduction/alkylation, followed by proteolytic digestion. These were merged into a single step in a smaller volume of 50 µL, which also increased the molar concentration of trypsin. At first, this reduced the number of peptides and proteins identified by 5% and 21% respectively, while also decreasing digestion efficiency (**Figure 4a**). Therefore, to compensate for the smaller reaction volume, the concentrations of reduction and alkylation reagents were optimised (**Figure 4b, V2**). Although this produced 30% more identified peptides and 7% more proteins, and carbamidomethyl-modified cysteine residues were unchanged, the digestion efficiency was poorer than for ABLE (**Figure 4b**). For example, 13% of the total identified peptides contained non-specific cleavages (up from 10% with ABLE). The relatively high number of non-specific cleavages suggested the presence of endogenous proteases other than trypsin/Lys-C in the FF samples. Therefore, a snap heating step was introduced at 95_C for 7 min in the presence of 5% SDC prior to sample lysis. This had a large impact on the proteins identified, 28% more peptides and 38% more proteins than ABLE, and improved digestion efficiency (reflected as fewer non-specific cleavages - from 18% to 2%) (**Figure 4c, V3**). The data suggests that endogenous protease activity released upon lysis was indeed a confounding issue in FF samples.

**Figure 4:**
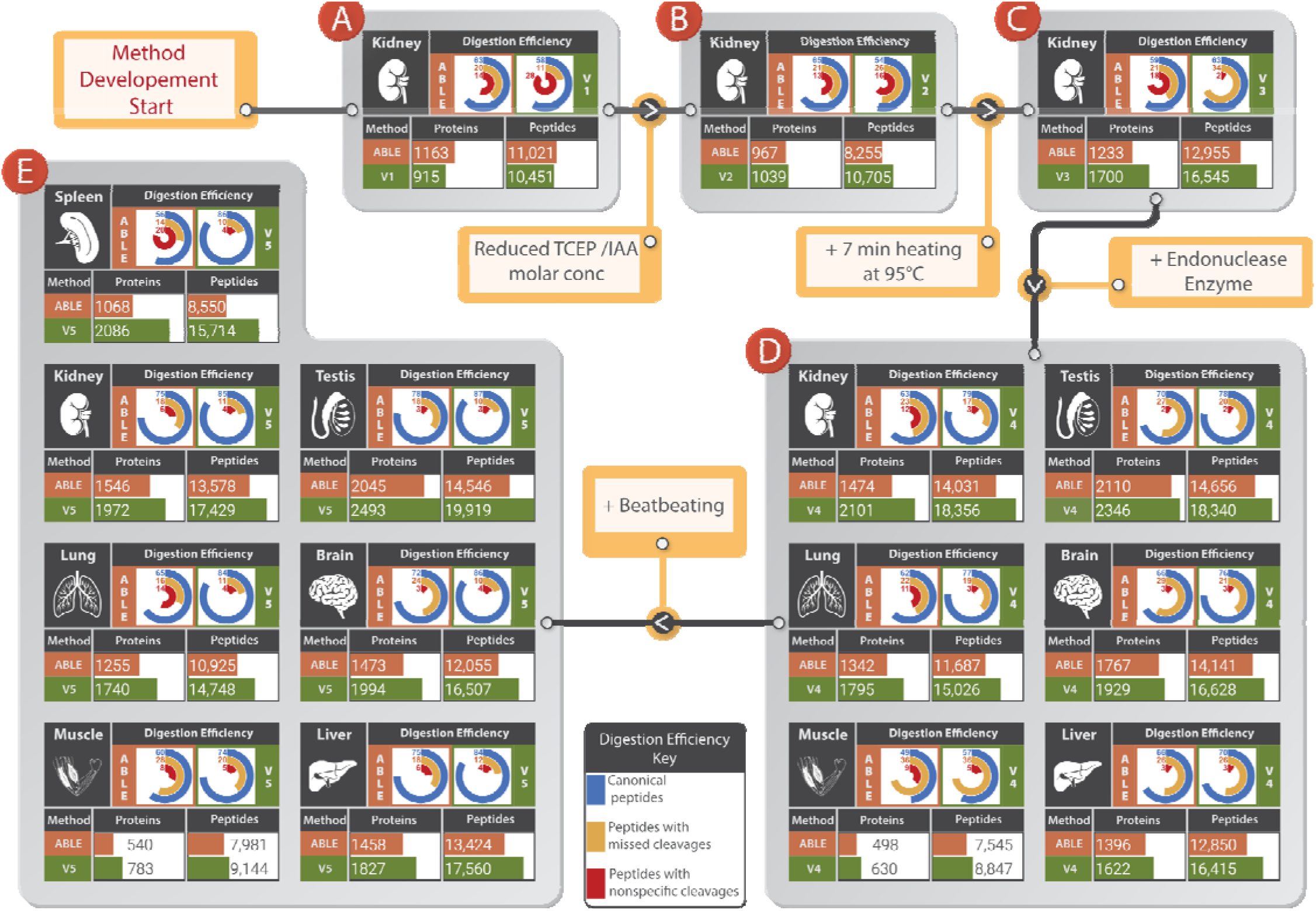
Heat ‘n Beat (HnB) method development in fresh frozen (FF) tissue punches. The HnB method development process involved 5 steps, each representing different iterations in the method development (V1-V5), each compared to the previous ABLE method. All tissues were collected as tissue punches from FF rat organs. Main modifications made to improve the preceding method (V1-V4) are indicated between each panel. **a**) compares ABLE to a single-step new V1 method. **b**) compares ABLE to a single-step V2 method with reduced TCEP and IAA concentrations to improve trypsin/Lys-C activity. **c**) compares ABLE to the new V3 method with an added heating step at 95°C to denature endogenous proteases to reduce protease activity and thereby reduce non-specific cleavage. **d**) compares ABLE to a new V4 method with the addition of Benzonase to cleave nucleic acids and aid trypsin/Lys-C digestion by reducing steric hindrance. **e**) compares ABLE to a the preceding method with introduction of a bead-beating step after heating and prior to digestion, V5, to more completely lyse tissue and further aid trypsin/Lys-C activity by reducing steric bulk. The panels list the tissue type/s used for testing. The mini-tables show the number of peptides and proteins identified, and the protein yields obtained using the given method. The radial bar graphs show the -/Lys-C digestion efficiency for each tissue section.

On the hypothesis that polymeric DNA and RNA could hinder access of trypsin/Lys-C to proteins, Benzonase nuclease, a promiscuous endonuclease, was introduced to cleave them and reduce sample viscosity in HnB V4 ^34^. For the FF kidney, this produced a further substantial increase in peptides and proteins identified by HnB by 31% and 43% respectively, while also reducing the number of missed and non-specific cleavages (**Figure 4D, V4**). The V4 method was next applied to punches from six FF tissues from multiple rat organs representing an expected range of homogenisation difficulty (kidney, muscle, liver, brain, testis and lung). The overall improvement was smaller for tissues that are more difficult to homogenise, including muscle and liver. We noted that such samples sometimes contained small pieces of intact residual tissue after the Barocycler, suggesting incomplete tissue disruption for these samples. Therefore, a final step was introduced for physical disruption of the samples with a bead-beater ^10,35,36^ in the presence of zirconium beads and SDC for 1 min, prior to the Barocycler step (**Figure 4E, V5**). This final change improved protein and peptide identification in these two ’difficult’ tissues, as well as substantially improving digestion efficiency in five of the seven tissues. Improved protein and peptide identifications were obtained in all seven FF tissues, with the greatest improvement of 95% and 85% respectively being in observed in spleen. Overall, the five cumulative substantial changes to the ABLE protocol comprise the final 38 min HnB protocol, performed in a single low volume Barocycler MicroTube.

### HnB performance across diverse samples

Having developed the HnB method with diverse FF solid tissue punches, we assessed performance of the protocols with the two different tissue preparation methods, FF-OCT embedding and FFPE fixation (both being cut as 10 µm sections) across six tissue types (n = 4). Two different washing or SPE methods as described above were used as appropriate for the relevant embedding or fixation method for the two cohorts. Overall, an across-the-board improvement was obtained for both FF-OCT and FFPE samples in the peptide yields, and number of peptide and protein identifications using the final HnB protocol (**Figure 5**). For example, the improvement for FF-OCT tissues was up to 85% and 95% more peptide and protein identifications respectively, accompanied by an increase in canonical peptides up to 79-89% compared with 65-80% using ABLE (**Figure 5a**). We attribute the improvements in peptide and protein identifications to the more complete protein digestion for almost all samples. Non-specific cleavage was most improved in FF and FF-OCT samples (**Figure 4e vs 5a**). As expected, there was less improvement in non-specific cleavages with FFPE tissue, presumably since the formaldehyde fixation likely resulted in less endogenous protease activity released upon homogenisation. Regardless of the sample source, brain tissue showed little change in non-specific cleavage. A higher proportion of canonical tryptic sequences in a digest increases the potential for search algorithms to accurately identify peptides and proteins while decreasing sample complexity and thereby reducing computational requirements and potential FDR challenges. HnB outperformed ABLE with every tissue type across each preservation technique (**Figure 4E & 5**). Specifically, 84-90% of peptides identified using HnB across most tissue types were canonical tryptic sequences, with muscle tissue the only exception, where 72-77% of the identified peptides were canonical. The number of canonical tryptic sequences identified using ABLE was much lower at 56-79% and dropped further to 53-65% for muscle tissue.

**Figure 5:**
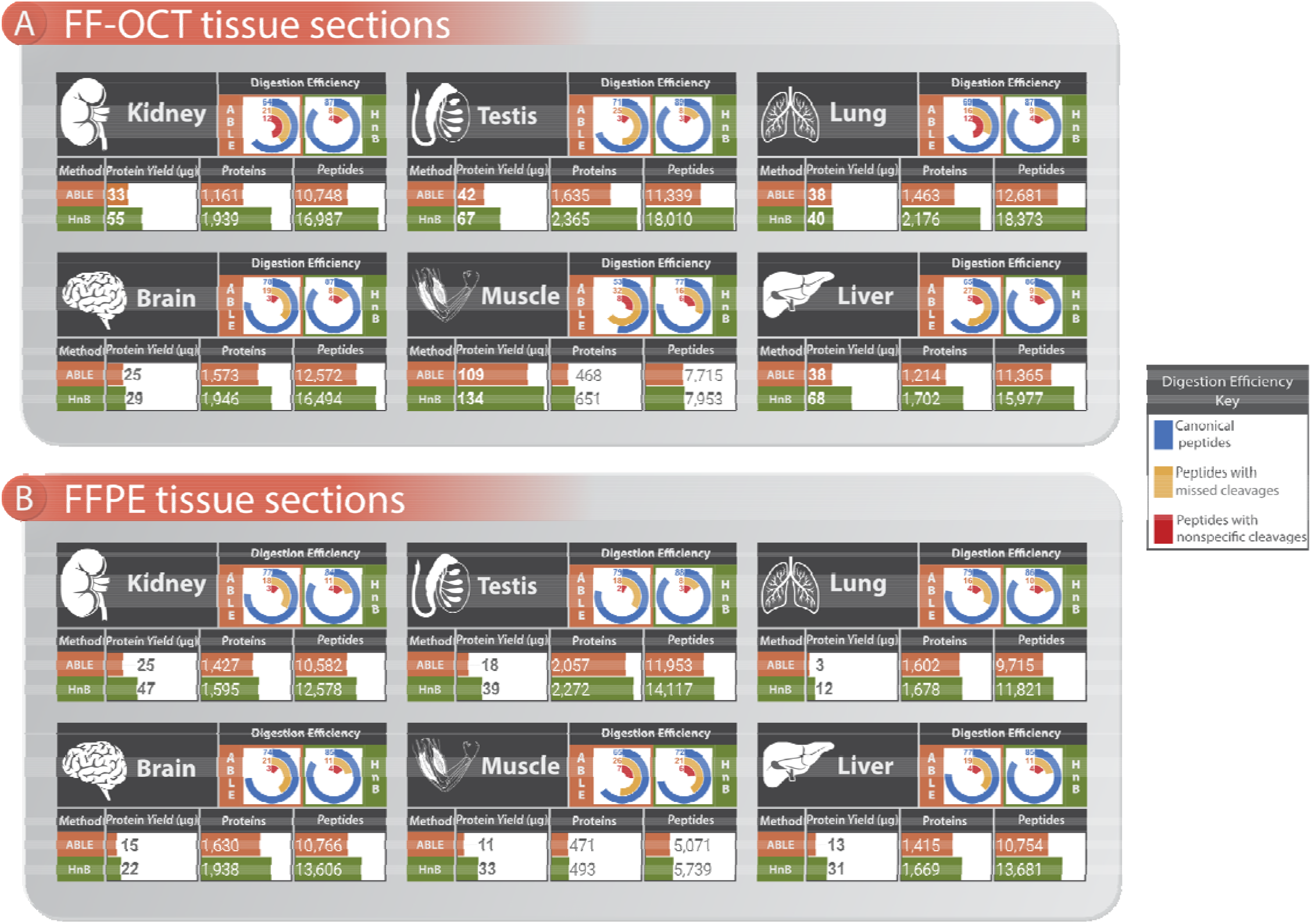
Application of HnB to FF-OCT or FFPE tissue sections. The HnB (V5 from **Figure 4e**) and ABLE methods were compared on tissue from six different rat organs, including 30 µm FF-OCT tissue sections and 10 µm FFPE tissue sections. Panel a) displays the results when 30 µm FF-OCT tissue sections were prepared using HnB or ABLE, while panel b) shows the results when 10 µm FFPE tissue sections were prepared using HnB or ABLE. The mini-tables show the number of peptides and proteins identified, and the protein yields obtained using the given method. The radial bar graphs show the trypsin/Lys-C digestion efficiency for each tissue section.

Another index of method improvement is the overall peptide yield, as greater yield equates to more sample for MS analysis, or to the requirement for less sample. The sectioning of FF-OCT or FFPE tissue blocks introduces a small variability in tissue volume between specimens due to orientation of the block between samples. Peptide yield increased across the entire range of samples with HnB (**Figure 5**). The largest improvements were achieved for FFPE tissues, with up to 390% increased peptide yield, while increases of up to 80% were obtained for FF-OCT tissues. We attribute the improved yield primarily to the bead-beating step.

### HnB identifies additional proteins

Samples prepared using HnB resulted in significantly more peptides and proteins being identified by MS than ABLE, with almost all tissue types across the different sample classes (FF-OCT embedded muscle and FFPE lung, whilst greater, were not significant) (the statistics are shown in **Supplementary Table S2**). The number of proteins identified improved by up to 95% for FF punches, by up to 75% for FF-OCT sections and up to 19% with FFPE sections (**Figures 4-5**). The proteins identified for the various FF-OCT tissue sections overlap around 50% (**Supplementary Figure S2**). However, HnB identified up to 48% unique (not seen with ABLE) proteins whereas ABLE identified up to 10% unique (not seen with HnB) proteins. This shows that HnB provides an extended proteome coverage with minimal loss compared to ABLE. When the comparison was applied to a FF human HEK293 cell line, the HnB protocol achieved a 30% and 26% improvement in identified peptides and proteins, accompanied by an increase in the number of canonical tryptic peptides (**Supplementary Figure S3**), indicating the improved performance of HnB also applies to cell line pellets.

### Comparison of different tissue embedding techniques

To determine how the three different types of samples compare in quantitative terms, DIA/SWATH MS acquisition was used. The overall protein overlap between the three methods ranged from 54.7% to 82.2% with the lowest being muscle and testis the highest. A slightly larger number of identifications were found in the FF sample (**Supplementary Figure S4a-e**). The protein identifications were analysed by principal component analysis (PCA) for 5 tissues (brain, kidney, liver, muscle and testis) and sample preservation types (FF, OCT and FFPE) (**Supplementary Figure S4f**). Identifications from all tissue types clustered well, independent of the three sample preservation techniques.

### HnB technical performance

While optimising HnB it was noted that the digestion efficiencies for several samples with excessively large tissue punches remained high, despite the resulting protein to enzyme ratio being much higher than the recommended ratio of 20:1. The effect of increasing quantities of FF rat liver tissue punches on digestion efficiency was examined, where the amount of proteolytic enzyme was unchanged. As the amount of tissue increased, digestion efficiency decreased, yet remained greater than 70% canonical tryptic peptides, despite the protein to enzyme ratio reaching 300:1 (**Supplementary Figure S5, Bead-beater 1**). Another bead-beater instrument (MP FastPrep-24 and Qiagen TissueLyser II) with a higher speed (6.5 m/s) further improved the digestion, with samples achieving over 80% of canonical tryptic even when the protein to enzyme ratio was greater than 500:1 (**Supplementary Figure S5, Bead-beater 2**). High digestion efficiency well beyond the recommended protein to enzyme ratio of 20:1 indicates that tissue size is an unlikely source of method variability while also substantiating HnB robustness.

The completeness of the reduction and alkylation processes was examined. Essentially complete modification of cysteine residues with either HnB or ABLE methods was achieved, but HnB samples contained a greater number of carbamidomethyl modified peptides than ABLE (**Supplementary Figure S6a**). The revised quantities of the reducing and alkylating reagents in HnB lowered off-target *N*-terminal alkylation (**Supplementary Figure S6b**).

### The role of the Barocycler

ABLE and HnB were initially enabled by Barocycler technology, but due to the addition of the upstream steps, notably bead-beating, we examined whether there remained a need to retain the barocycling step in the final HnB method. Without a Barocycler, HnB showed a small non-significant reduction in peptides (*p*=0.053), but not in protein identifications, and significantly lowered the digestion efficiency (*p*= 0.0010) for rat kidney tissue (**Supplementary Figure S7**). Overall, Barocycler-assisted digestion still presents an advantage, even with an incorporated bead-beating step, since improved digestion efficiency will potentially translate to better reproducibility over time, especially for hard to lyse tissues.

The collective data indicates HnB is a robust method with significant quality improvement across multiple sample types and embedding and fixation methods. It requires substantially less handling due to the removal of all tube changes prior to SPE and results in significant improvements in time saving, peptide quality and quantity.

### Application to large scale/high-throughput cohort

The suitability of HnB was evaluated for clinical samples using eight FF-OCT embedded human prostate tumour biological replicate punches. The HnB protocol identified 62% more peptides and 81% proteins than ABLE in prostate samples (**Supplementary Figure S8**). Digestion efficiency also increased to 85% of peptides identified as canonical as opposed to 64% with ABLE. Consequently, the method appears to have broad applicability across tissue types.

To determine how HnB supports a high-throughput workflow 1,171 patient samples from 25 different tissue types were processed with HnB from FF-OCT sections (**Figure 6**). For most organs, the cohort comprised tumour and normal tissues from the same organ. The clinical outcomes of this study will be reported elsewhere, here we report the technical overview. The sample core processing time was 60 hours, with a further 32 hours required for MCX SPE clean-up (92 hours total sample preparation time). The samples were run in triplicate across six 6600 TripleTOF mass spectrometers in DIA/SWATH mode, comprising 3,513 runs and including standards brings this to 4,083 LC-MS runs of 90 min each, totalling 6,124 hours of acquisition time (8.5 months if a single mass spectrometer was used). The acquisition time can be greatly reduced with the use of multiple instruments in parallel and shorter run times. In most tissue types approximately 4,000-6,000 quantifiable proteins were identified (**Figure 6c**). Tissue with the highest fat content: including adipose tissue (both tumour and normal), soft tissue not otherwise specified (NOS) and normal breast tissues produced lower protein identifications. Bone and muscle were also lower. The digestions were consistently of excellent quality; with peptide yields all being high, ranging from 27-80 µg of peptides (**Figure 6c**) and digestion efficiencies were greater than 80%, illustrated with the mis-cleavage data of around 15% for most tissues (**Figure 6d**). Using conventional OCT washing methods and overnight digestion methods (2 hr wash plus 20 hr core processing, per sample) this cohort could have taken up to 77 days (∼2.5 months) for sample preparation alone (in batches of 15). Overall, this shows remarkable throughput and output performance for HnB. The data demonstrates the large impact of this end-to-end method in accelerating high-throughput proteomics.

**Figure 6:**
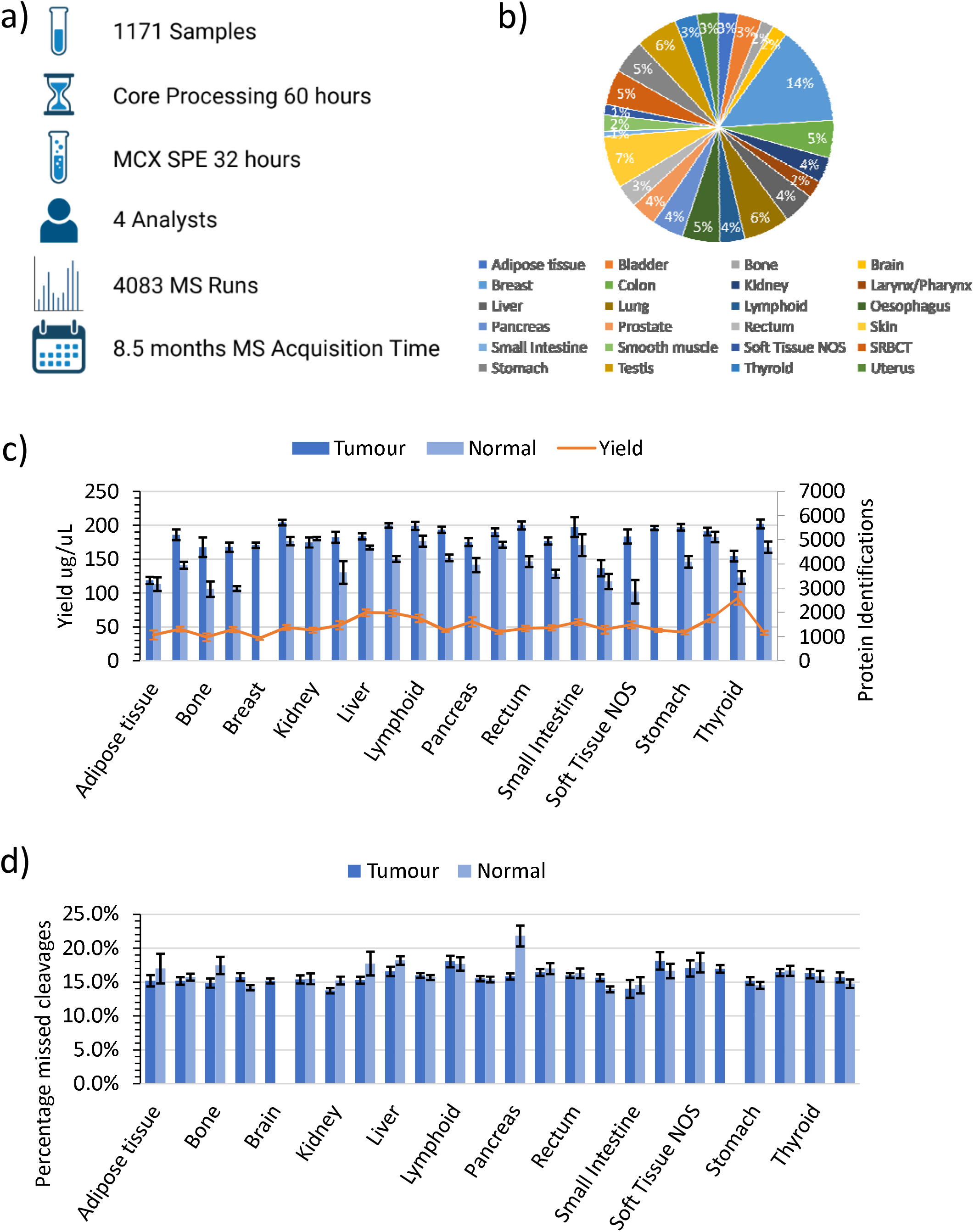
Application of HnB to a large scale cohort of human cancer biopsies. **a)** Summary of processing and MS time for a library of 1,171 human cancer samples. **b)** Distribution of the 1,171 human samples between 25 tissue types. **c)** Peptide yield and number of protein identifications following MS, using the final HnB protocol (V5 from **Figure 4e**). All data are means ± SEM for n = 3 technical replicates **d)** Digestion efficiency is displayed as miscleavages per tissue type.

## Conclusions

In this report each step of the MS sample preparation process was systematically tested and improved. Initially rat kidney tissue was used for method development, then the methods were expanded to six other rat tissues and cell lines for breadth of applicability, and finally to a large biobanked human tissue cohort for expandability. Key amendments made to the original sample preparation method include the addition of a new heating step to reduce endogenous protease activity, a bead-beating step to improve the efficiency of tissue homogenisation and lysis, addition of an endonuclease to improve trypsin/Lys-C digestion efficacy and modifying the reducing and alkylating conditions to promote trypsin/Lys-C enzyme activity and reduce off target alkylation. A critical development was our demonstration that the time-consuming OCT removal procedure is not required at the start of the method for FF-OCT samples, but can be shifted to the final SPE step, potentially saving hours for this class of sample. Combined with the development of an MCX-OCT clean-up protocol and optimisation of the FFPE methanol-heptane deparaffinisation protocol, we have produced an overall streamlined end-to-end protocol that is simple to implement, robust, and which yields high quality tryptic peptides in under an hour from diverse tissue samples.

Traditionally, FF, FF-OCT and FFPE tissues require different sample preparation pipelines, that limits the ability to identify biomarker trends across preservation techniques. There can be anywhere between 47-92% overlap of protein identification between FF and FFPE tissues ^32,37,38^. Previous studies have sought to optimise individual steps in the sample preparation workflow, notably the deparaffinisation or the tissue solubilisation steps for FFPE samples ^12,39^, the LC settings ^40^ or the type of MS fragmentation ^15^. However, to the best of our knowledge none have taken an end-to-end approach or incorporated such a breadth of tissue sources. By identifying the limitations of existing methodologies and adapting them to suit various tissues, HnB was developed as a one tube, universal sample preparation method for proteomic analysis by MS. The resulting method delivered samples that yielded greater protein recoveries, better proteome coverage with more proteins identified and improved reproducibility all whilst reducing sample preparation time. Together with improvements of the FFPE dewaxing protocol and the development of the MCX protocol which eliminates the laborious task of tissue washing to remove OCT, a streamlined overall protocol has been developed that can be used to process the vast majority of clinically available samples in the shortest possible time while obtaining an improved proteome coverage.

The use of HnB for FF-OCT or FFPE samples is likely to have key clinical relevance in the future for fast sample analysis to support diagnosis and biomarker identification in patients. Applying this method to a large clinical cohort allowed the end-to-end preparation and processing of 1,171 various tumours for MS in just 92 hours, with data acquisition in triplicate being dependent on the gradient length and instrument model, but which in our hands took an 8.5 month period averaged for a single 6600 TripleTOF. This cohort included difficult to solubilise tissues such as smooth muscle and skin, with consistent reproducible results across all types and low missed cleavage rates between 15-20%. One of the main benefits for this cohort was the use of MCX clean-up for the removal of OCT, which saved many hours of labour-intensive washing and sample loss. Combining rapid sample preparation time and short MS runs, using faster MS instrumentation, illustrates the potential of this method in clinical pathology in the future.

## Acknowledgements

The authors wish to thank Peter J. Wild (University Hospital Zurich, Switzerland) for supplying prostate tissue. This work was supported by the Australian Cancer Research Foundation, Cancer Institute New South Wales (NSW) (2017/TPG001, REG171150), NSW Ministry of Health (CMP-01), The University of Sydney, Cancer Council NSW (IG 18-01), Ian Potter Foundation, the Medical Research Futures Fund (MRFF-PD), National Health and Medical Research Council (NHMRC) of Australia European Union grant (GNT1170739, a companion grant to support the European Commission’s Horizon 2020 Program, H2020-SC1-DTH-2018-1, ’iPC – individualized Paediatric Cure’ [ref. 826121]), and National Breast Cancer Foundation (IIRS-18-164). The work at ProCan^®^ was done under the auspices of a Memorandum of Understanding between Children’s Medical Research Institute and the U.S. National Cancer Institute’s International Cancer Proteogenomics Consortium (ICPC), that encourages cooperation among institutions and nations in proteogenomic cancer research in which datasets are made available to the public. P.J.R. is supported by an NHMRC Fellowship (GNT1137064). The Victorian Cancer Biobank through the Cancer Council Victoria as Lead Agency is supported by the Victorian Government through the Victorian Cancer Agency-a business unit of the Department of Health and Human Services.

## Author contributions

Conceptualization: DX, NL, PGH, PJR

Methodology: DX, NL, SGW, JMSK, CL, PGH

Software: NL, KA

Formal analysis: DX, NL

Investigation: DX, NL, SGW, JMSK, KA, PGH

Data curation: DX

Project administration: PGH, RR, PGH

Writing – original draft: NL, DX, PGH, PJR

Writing – review & editing: All authors.

## Supplementary Tables and Figure Legends

**Supplementary Table S1.**
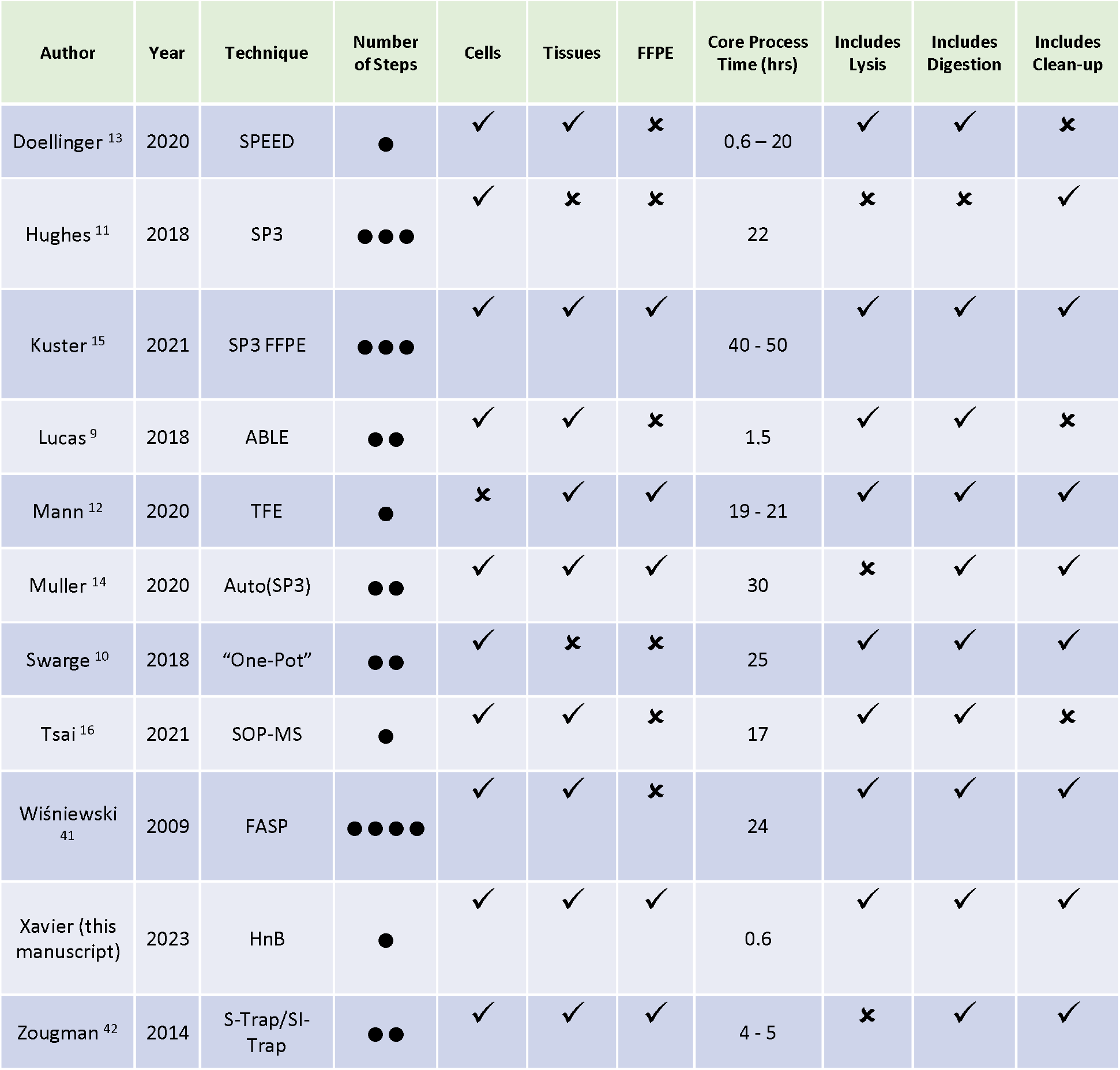
Examples of current high throughput proteomics methods.

**Supplementary Table S2.** Statistical data for HnB identifying more proteins than ABLE. *p*-values for peptides and proteins identified by MS across all rat organ types and across the different sample classes observed to differ between HnB and ABLE.

|  | *p*-value<br>Peptide | *p*-value<br>Protein |
| --- | --- | --- |
| Muscle FF | 0.049575 | 0.002925 |
| Muscle FF-OCT | 0.134662 | 0.000190 |
| Muscle FFPE | 0.000066 | 0.029432 |
| Liver FF | 0.000387 | 0.000998 |
| Liver FF-OCT | 0.000003 | 0.000027 |
| Liver FFPE | 0.000003 | 0.000215 |
| Lung FF | 0.000071 | 0.000418 |
| Lung FF-OCT | 0.000006 | 0.000002 |
| Lung FFPE | 0.000013 | 0.050499 |
| Kidney FF | 0.000024 | 0.000026 |
| Kidney FF-OCT | 0.000000 | 0.000001 |
| Kidney FFPE | 0.000045 | 0.000486 |
| Brain FF | 0.000005 | 0.000014 |
| Brain FF-OCT | 0.000006 | 0.000069 |
| Brain FFPE | 0.000030 | 0.000023 |
| Testis FF | 0.002261 | 0.011013 |
| Testis FF-OCT | 0.000066 | 0.000544 |
| Testis FFPE | 0.000078 | 0.000174 |
| Spleen FF | 0.000002 | 0.000004 |

**Supplementary Figure S1.**
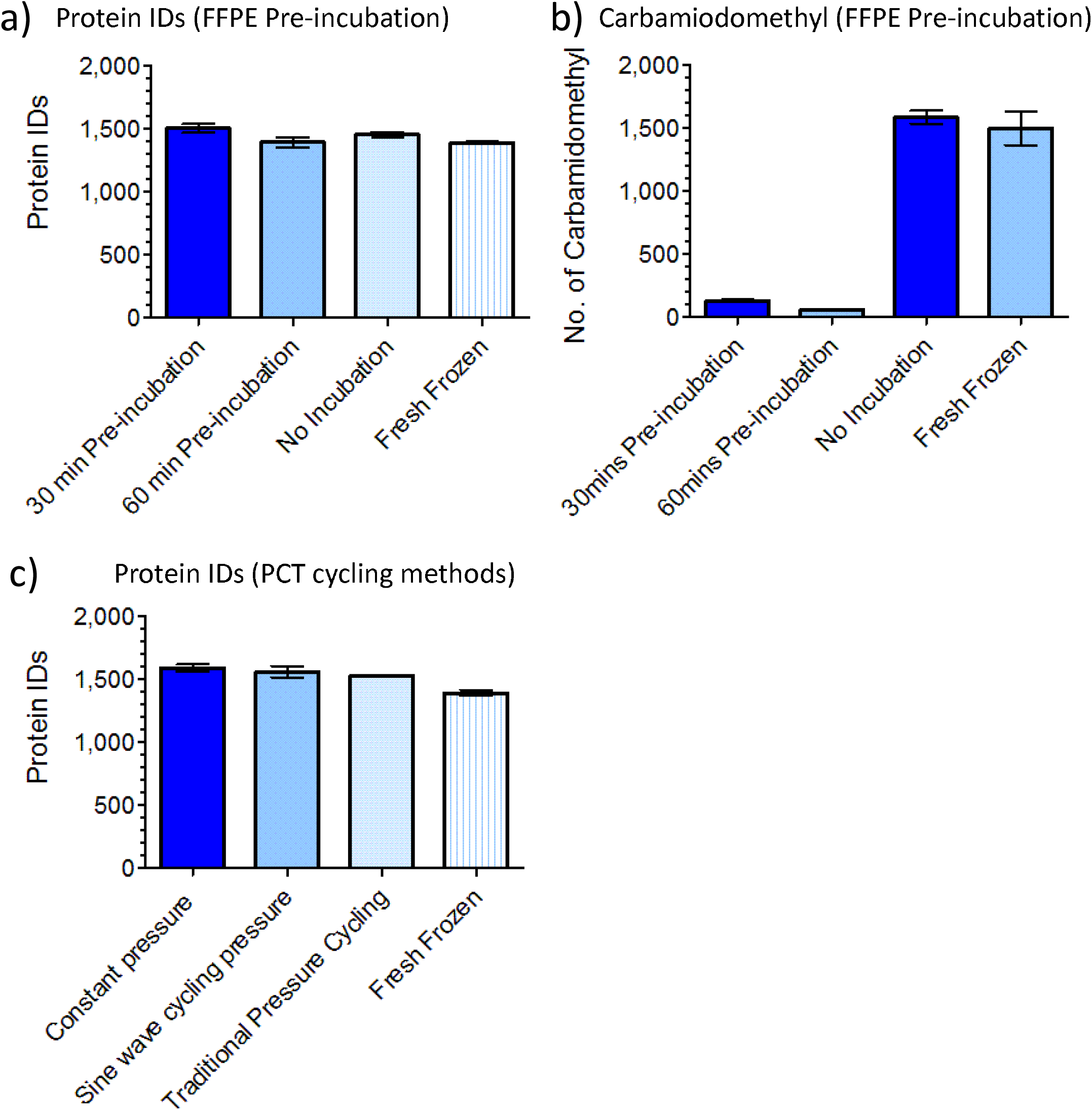
A pre-incubation step, occurring prior to tissue lysis, was tested on rat kidney FFPE sections. Samples were incubated at 100_C for either 30 or 60 min. Lysis and digestion were then kept the same for all samples. The pre-incubation was compared to FFPE with no incubation and FF (also no incubation). Data are means ± SEM for n = 4 biological replicates. **a)** Number of average protein identifications. **b)** Number of modified carbamidomethyl groups. **c)** Effect of different types of pressure cycling on tissue lysis. Traditional PCT on a Barocycler involves applying pressure for 50 seconds and then back to ambient pressure for 10 seconds. This was compared with a constant pressure of 45 kpsi or a sine wave pressure which of 45 kpsi ± 5 kpsi. No significant advantage was found with any of the methods.

**Supplementary Figure S2.**
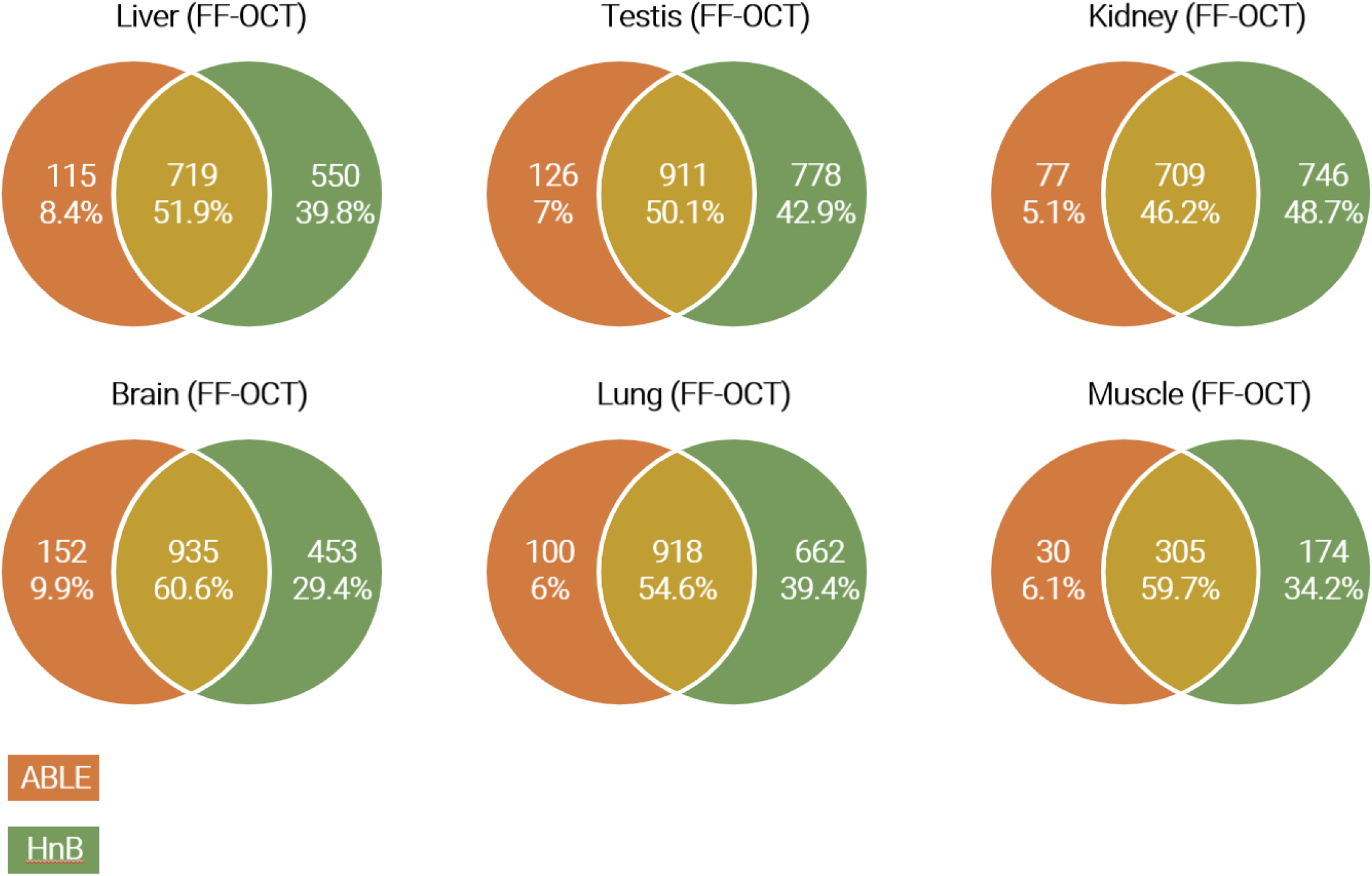
Overlap between proteins identified at 1% FDR when six tissue types preserved as FF-OCT were prepared using HnB and ABLE. Only proteins identified in all replicates for each method were used for the comparison.

**Supplementary Figure S3.**
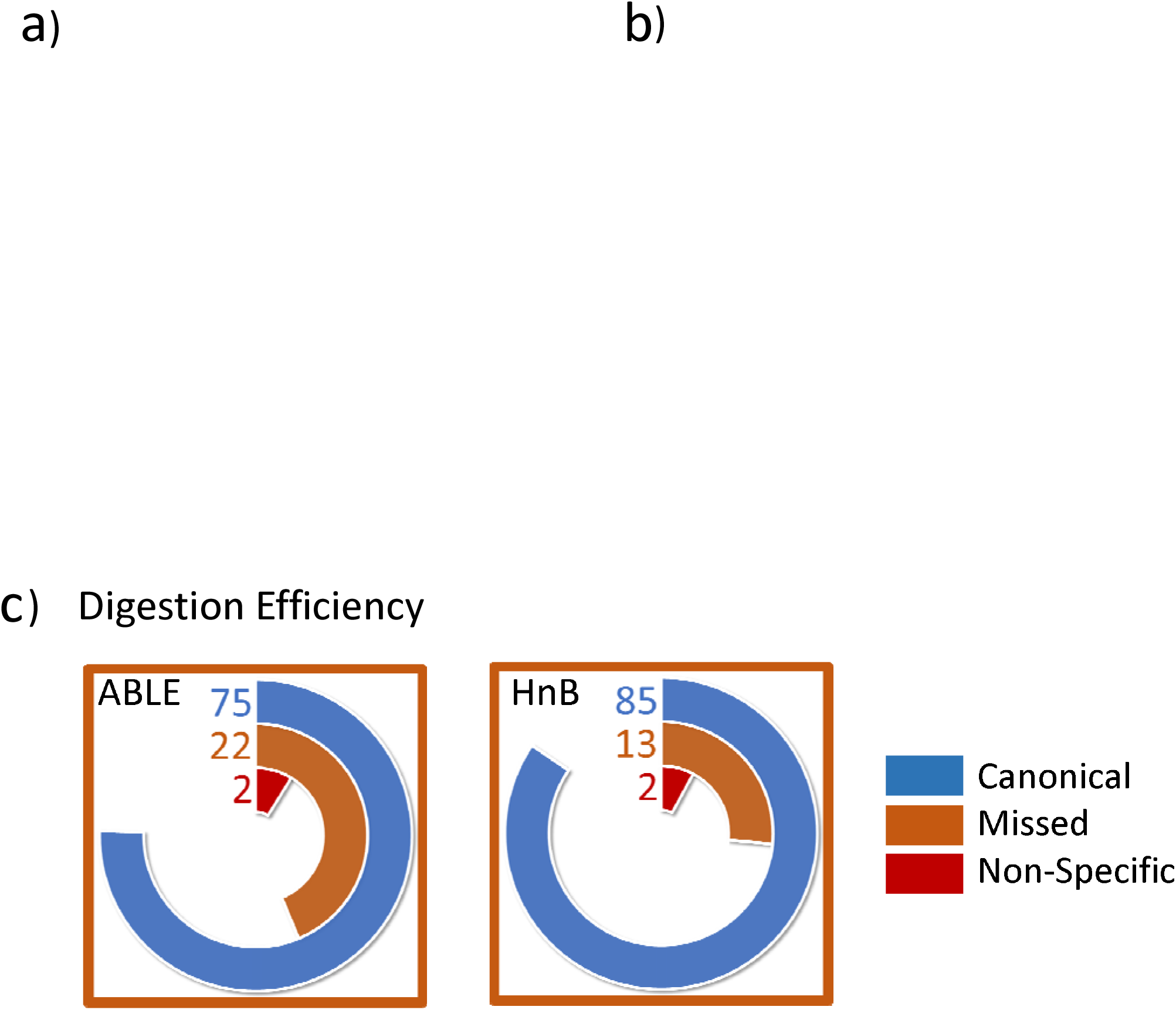
MS analysis of HEK293 cell lines processed using HnB or ABLE. **a**) Number of peptides identified at 1% FDR. **b**) Number of proteins identified at a 1% FDR. **c**) Digestion efficiency for HEK 293 cell lines when prepared using ABLE or HnB.

**Supplementary Figure S4.**
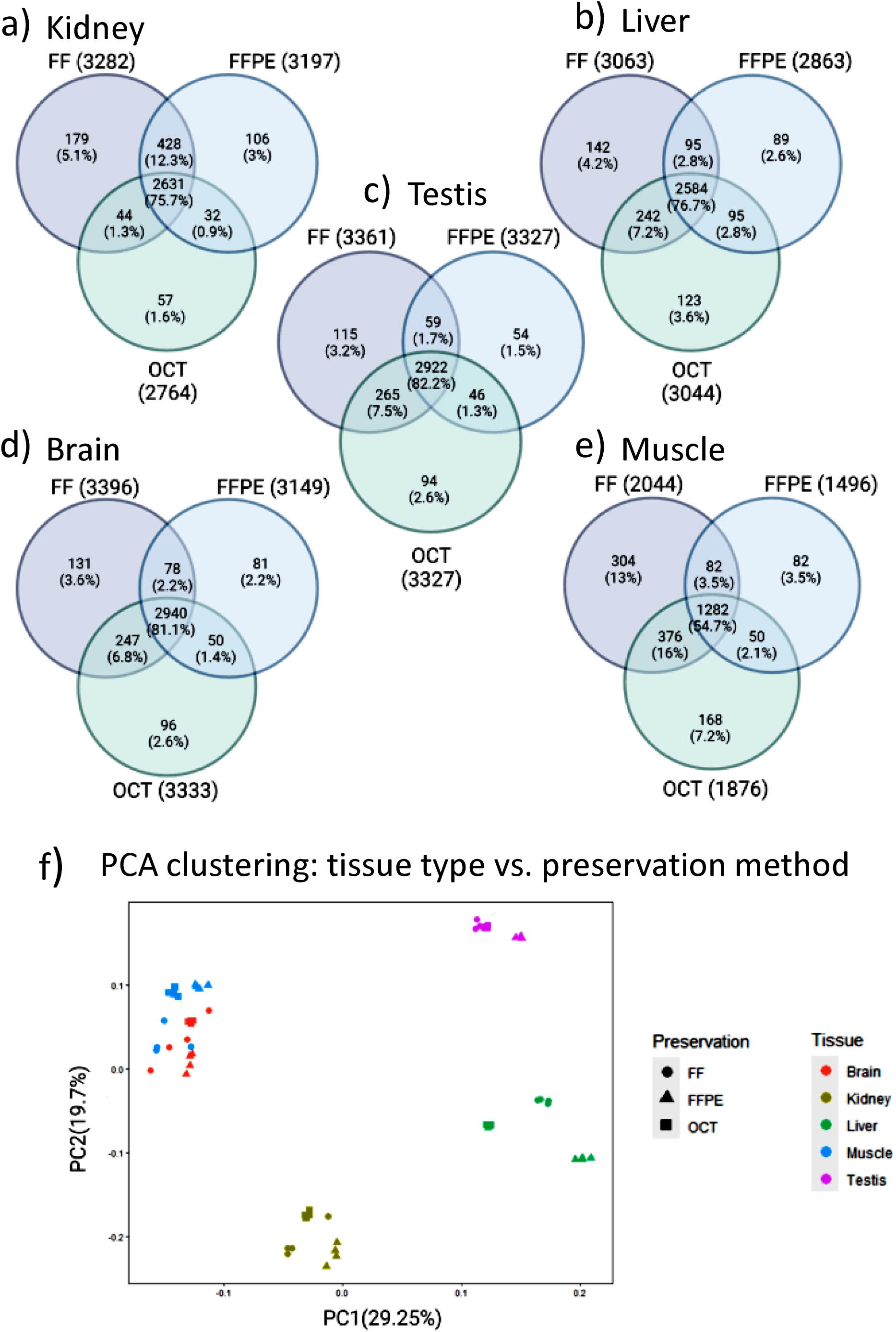
PCA comparing the different tissue types and preservation methods from DIA/SWATH MS data processed using DIA-NN software.

**Supplementary Figure S5.**
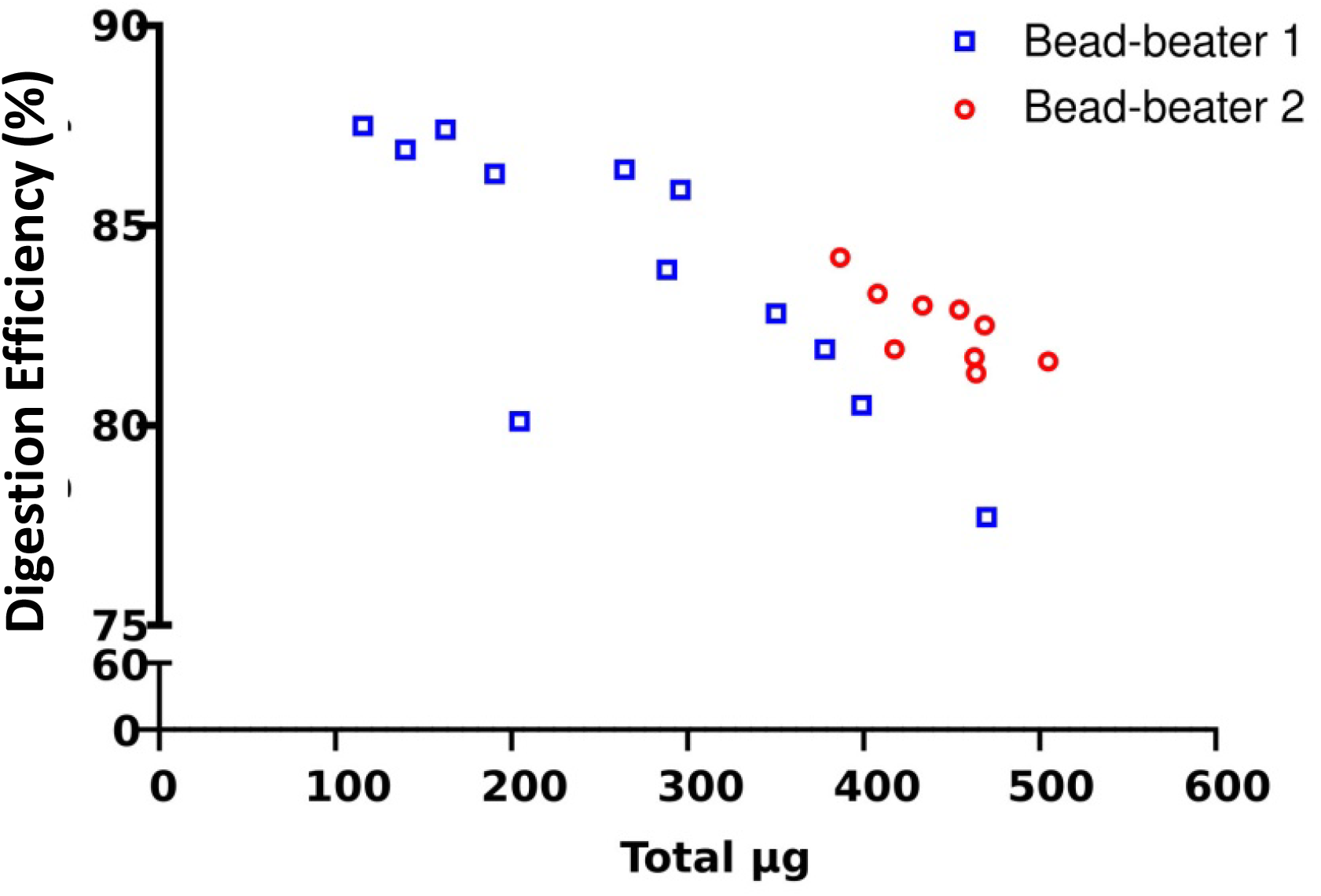
The effect of increasing quantities of FF rat liver tissue punches on digestion efficiency. Total percentage of canonical identified peptides for the given rat liver samples as the amount of tissue added to tubes was increased was used to determine its effect on trypsin/Lys-C digestion efficiency. Bead-beater 1 was a modified BeadBug™ microtube homogenizer while Bead-beater 2 was a Fastprep 24 classic.

**Supplementary Figure S6.**
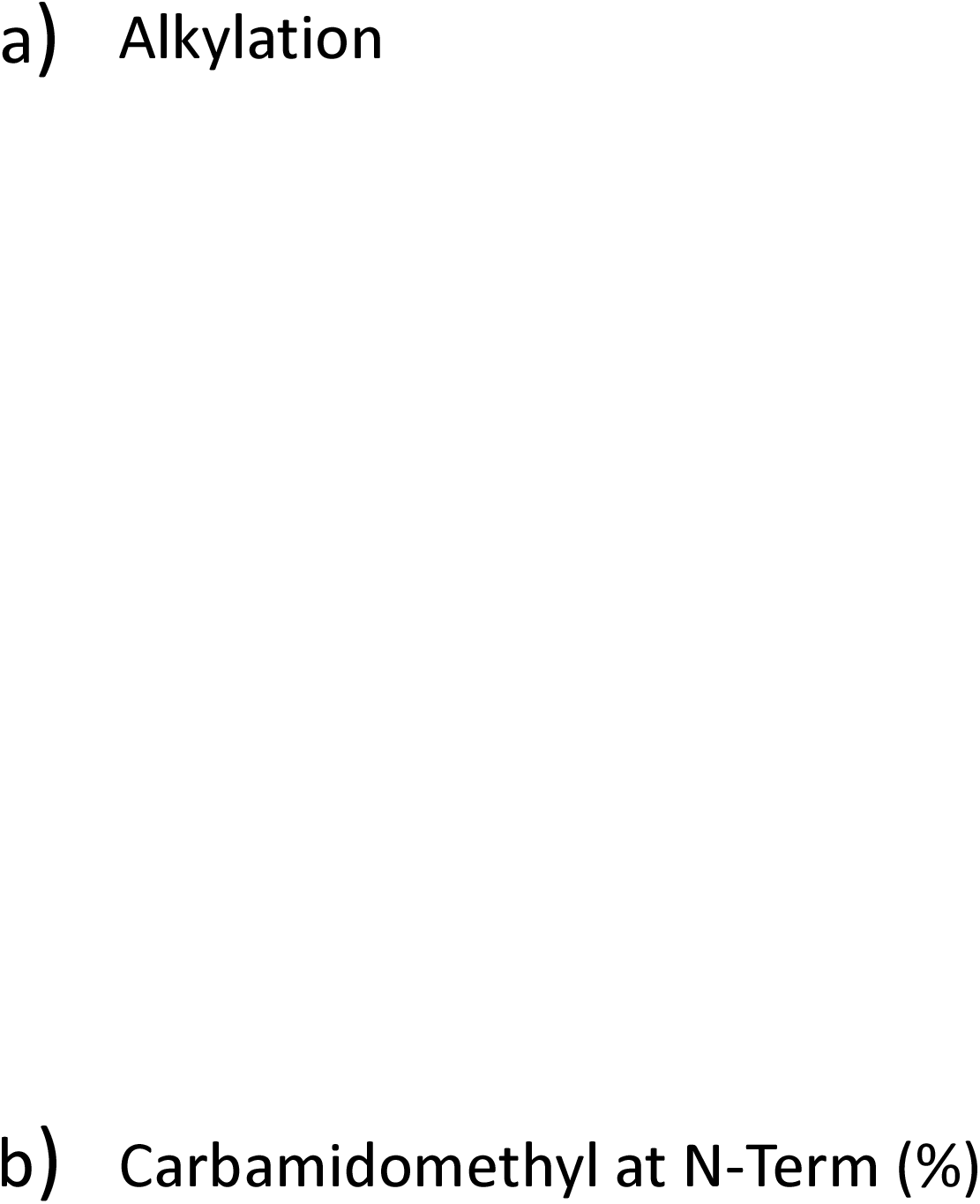
**a)** Number of identified peptides containing cysteine with the carbamidomethylation (CAM) modification when rat testis and rat liver which were preserved as FF-OCT and FFPE then prepared using HnB and ABLE. **b)** Percentage of total identified peptides containing *N*-terminal alkylation when rat testis and rat liver which were preserved as FF-OCT and FFPE then prepared using HnB and ABLE.

**Supplementary Figure S7.**
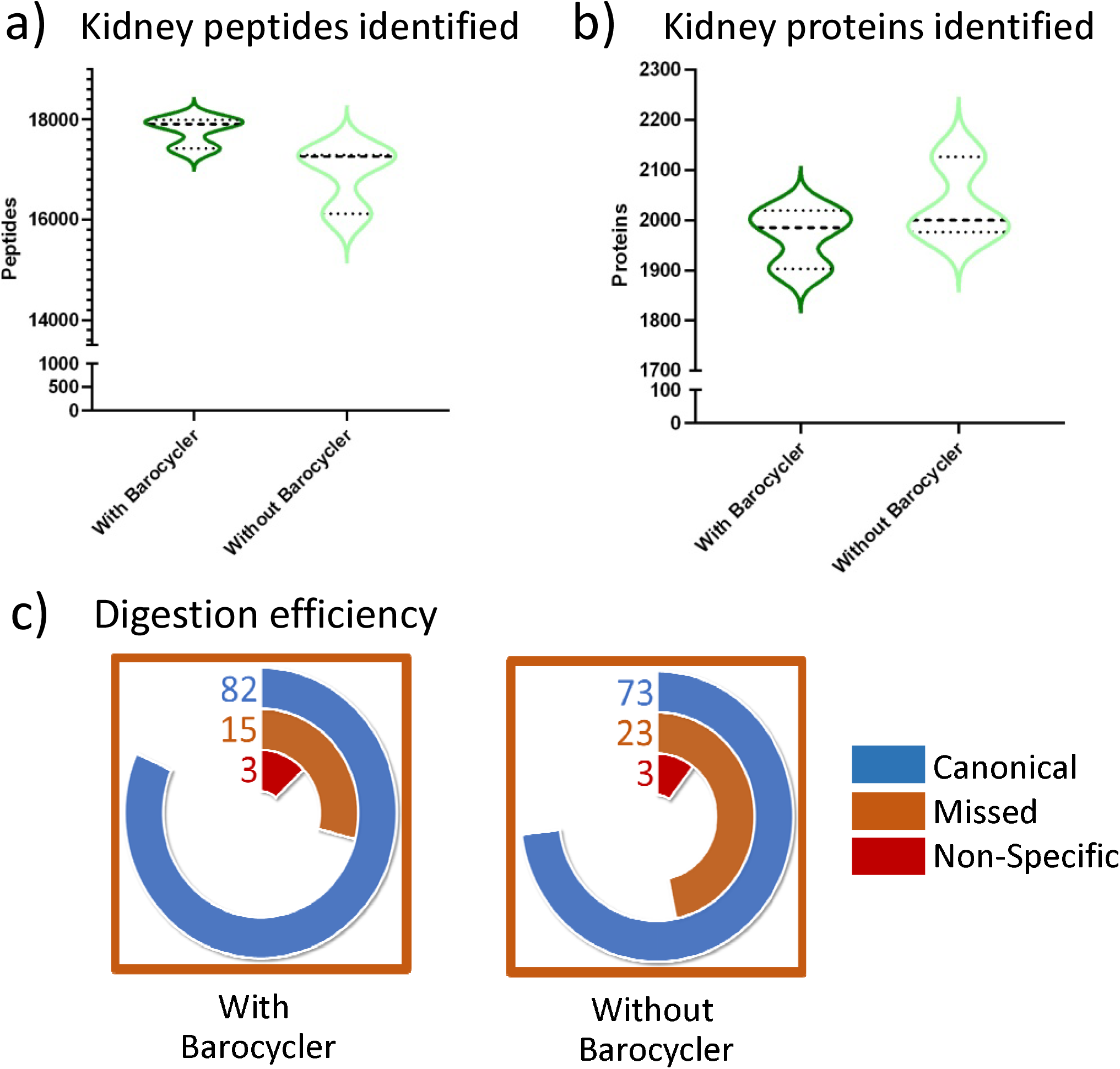
Role of the Barocycler in the HnB protocol. Results of FF rat kidney tissue punches prepared using HnB with and without a Barocycler. Samples not prepared in the Barocycler were placed in a water bath for 30 min at 70°C and compared to samples digested with trypsin/Lys-C in a Barocycler at 70°C without any pressure-cycling. **a**) Number of peptides identified. **b**) Number of proteins identified. **c**) Digestion efficiency expressed as a percent of canonical tryptic peptides, missed cleavage or non-specific cleavage. Paired t-tests: *p*=0.0526 (peptides); *p*=0.296 (protein); *p*= 0.0010 (digestion efficiency).

**Supplementary Figure S8.**
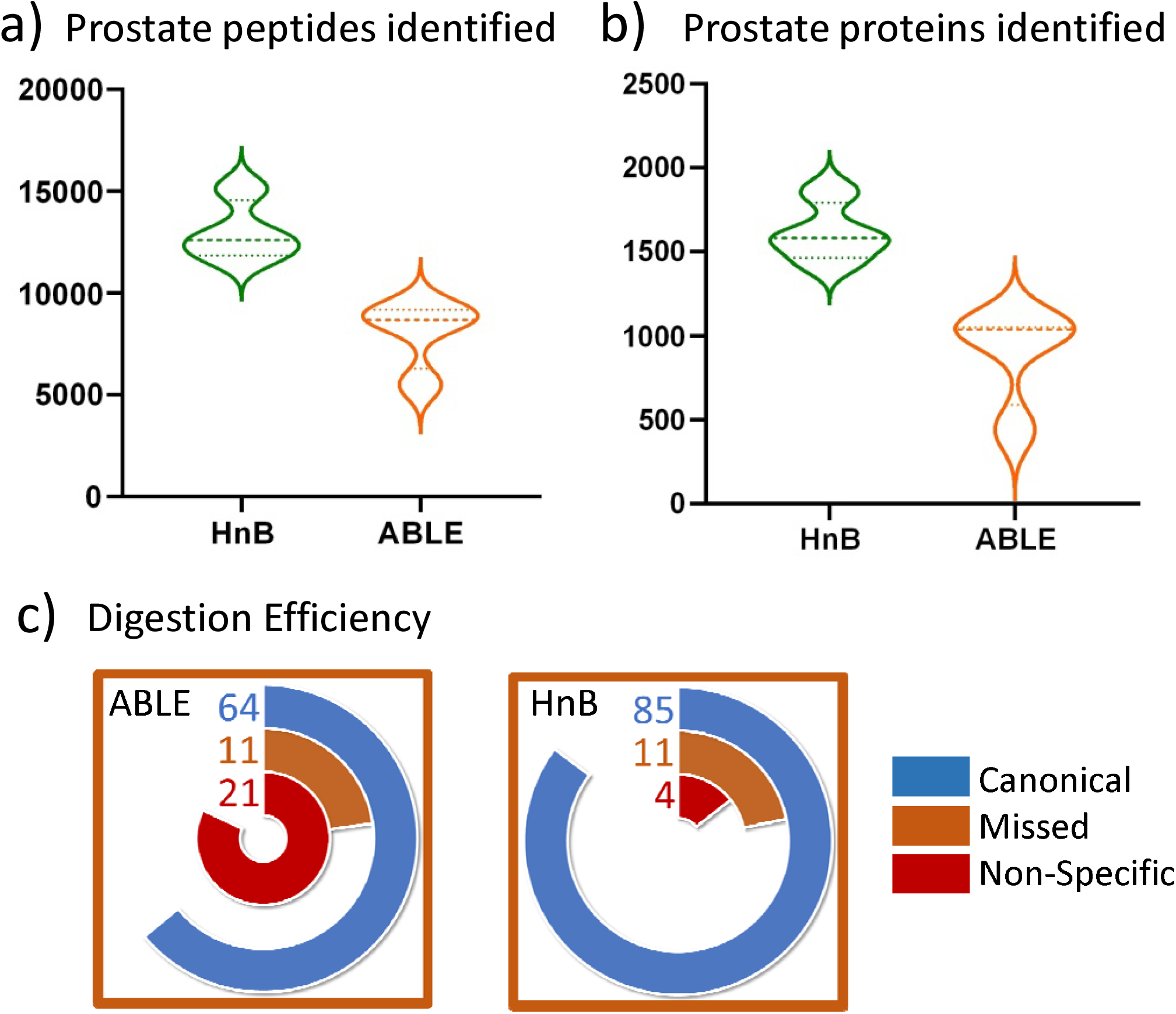
Application of HnB to prostate cancer sample. Mass spectrometry results when FF-OCT embedded human prostate tumour punches were processed using HnB compared with ABLE. **a)** Number of peptides identified at a 1% FDR was significantly better with HnB (*p* = 0.0039). **b**) Number of proteins identified at a 1% FDR was significantly better with HnB (*p* = 0.0080). **c**) Digestion efficiency expressed as a percent of canonical tryptic peptides, missed cleavage or non-specific cleavage.

